# Spatially heterogeneous inhibition projects sequential activity onto unique neural subspaces

**DOI:** 10.1101/2023.09.15.557865

**Authors:** Andrew B. Lehr, Arvind Kumar, Christian Tetzlaff

## Abstract

Neural activity in the brain traces sequential trajectories on low dimensional subspaces. For flexible behavior, these neural subspaces must be manipulated and reoriented within short timescales of tens of milliseconds. Using mathematical analysis and simulation of a recurrently connected neural circuit for sequence generation, we report that incorporating a subtype of interneurons that provides spatially heterogeneous inhibition enables the projection of sequential activity onto task- or context-specific neural subspaces. Depending on the sparsity of inhibitory projections, neural subspaces could be arbitrarily rotated, without altering the key aspects of sequence generation. Thus, we propose a circuit motif by which inhibitory interneurons can enable flexible switching between neural subspaces on a fast timescale of milliseconds, controlled by top down signals.

## Introduction

Sequential activity of neurons has been observed throughout the central nervous system as a reliable neural correlate of behavior, for example, in spinal circuits ^1^ and motor areas during movement ^2^, in higher cortical areas during decision making ^3^, and within the hippocampus during navigation and memory tasks ^4–6^.

In neural networks, sequential activity has been shown to live on low dimensional neural subspaces, which arise due to the correlation structure of the participating neurons (for review, see e.g. ^7–10^). Moreover, different tasks or behaviors are represented as a change in the orientation of the neural subspaces ^2,11–13^. Computational studies have suggested that reorienting the neural subspace generated by a recurrently connected network requires either incremental learning over days ^14^ or full rewiring of the network ^15^. Thus, it remains unclear as to how neural subspaces can be flexibly and dynamically reoriented on fast, behaviorally important time scales, in particular without disturbing the underlying sequential dynamics.

There are several ways to generate sequences. In randomly connected networks, sequences may arise due to the presence of particular connectivity motifs ^1,16,^ or via structure imposed by supervised or unsupervised learning ^18–22^. Alternatively, network structures such as discrete assemblies connected in feedforward chains ^23–25^ or continuous models with local connectivity ^26–35^ akin to distance-dependent connection probabilities observed in the brain ^36–38^ generate sequences via asymmetry in the local connectivity ^27,28,34^ or short-term neuronal or synaptic adaptation/depression ^29,31,32,35,39^.

In this setting, the functional role of inhibitory interneurons is to balance the recurrent excitation. A number of studies have observed broad non-specific inhibition ^40–42^ consistent with the role of interneurons in modulating gain ^43^ and providing a global “blanket” of inhibition upholding excitation-inhibition (EI) balance ^44–47^. However, given the diversity of interneurons in the brain ^48–54^, while certain interneuron subtypes may indeed play a role in maintaining the EI balance, it is likely that others play a different role.

Indeed, a number of studies have highlighted potential complimentary roles of different inhibitory neu-ron subtypes ^55–61^. For instance, inhibitory plasticity can play a stabilizing role in sequence-generating circuits ^62^. And instead of a global “blanket”, in some instances inhibitory connectivity can be non-random, with an overabundance of certain motifs ^63–65^. Inhibitory synapses can be spatially clustered ^66^, selectively connect to specific dendritic branch types ^67^, and coordinate their plasticity with excitatory synapses in close spatial proximity ^68^. Such structured connectivity could support more sophisticated computational functions beyond maintaining a global EI balance.

Notably the projection patterns of certain inhibitory subtypes can support spatially heterogeneous inhibition, instead of just a global “blanket” of inhibition. For example in the cortex, the narrow axonal projections of vasoactive intestinal peptide (VIP) interneurons inhibit nearby somatostatin (SOM) interneurons, thus enabling transient disinhibition of local pyramidal neurons, effectively making “holes in the inhibitory blanket” ^69,70^. Similar heterogeneously distributed inhibition can be observed in the olfactory bulb, where stimulus-specific spatial patterns of GABA release result from the projection patterns of local interneurons ^71,72^. Constrained by this spatial landscape or “blanket with holes”, a key signature of information processing across these regions is that neural activity unfolds across time not as static spatial patterns, but as spatiotemporal activity sequences or traveling waves, both in cortex ^34,73^ and in the olfactory bulb ^74^.

But the question remains, what role do these heterogeneous and stimulus-specific inhibitory patterns play in network dynamics and neural computation? Here we focus on a potential functional role of this interneuron diversity in influencing sequential neural activity to enable fast, dynamic, and flexible control of the low dimensional neural manifold. In both rate-based and spiking neural network models with spatially asymmetric local connectivity, we show that spatially heterogeneous, clustered inhibition can rapidly manipulate the neural subspace while preserving sequence generation. Thus, in this circuit, one inhibitory population is crucial for balancing excitation during sequence propagation and a second inhibitory population — which we refer to as selective inhibition — can form the necessary sub-network to shape the network activity, selecting unique neural subspaces and, by this, endows the network with the computational benefit of sequence selection on behaviorally relevant timescales. Thus, we propose a biologically plausible mechanism in which different interneuron subtypes perform complementary roles in storing and dynamically selecting between activity sequences in one recurrent circuit.

## Results

Neural activity of a subset of *N* neurons recorded for *T* time steps can be described as a matrix *A* ∈ ℝ^*N* x*T*^ with each column of the matrix containing the activity vector, e.g. firing rates, *r*_*t*_ ∈ ℝ^*N*^ at time *t*. A number of studies have reported that neural dynamics, though they exist in an *N* dimensional space (one axis per neuron), tend to be of much lower dimensionality see e.g. ^7,8,^. For example, applying principal component analysis to the activity matrix *A* from either neural data or a mathematical model for sequence generation reveals a *K* dimensional subspace with *K << N*.

Here we investigate how this low dimensional subspace can be dynamically manipulated on a fast timescale of tens of milliseconds. Starting with a recurrently connected network of excitatory and inhibitory neurons and a connectivity structure that enables sequence generation (Fig. 1a), we consider a second population of interneurons that provides spatially heterogeneous, clustered inhibition. By this we mean that a coactive ensemble of interneurons cluster their synapses onto a subset of the excitatory and inhibitory neurons responsible for sequence generation, with the remaining neurons receiving negligible input. We parameterize the sparsity of the selective inhibition in terms of the fraction of neurons silenced, *p*_*inh*_ ∈ [0, 1], and throughout this paper we explore the effect of this selective silencing on the low dimensional subspace.

**Figure 1.**
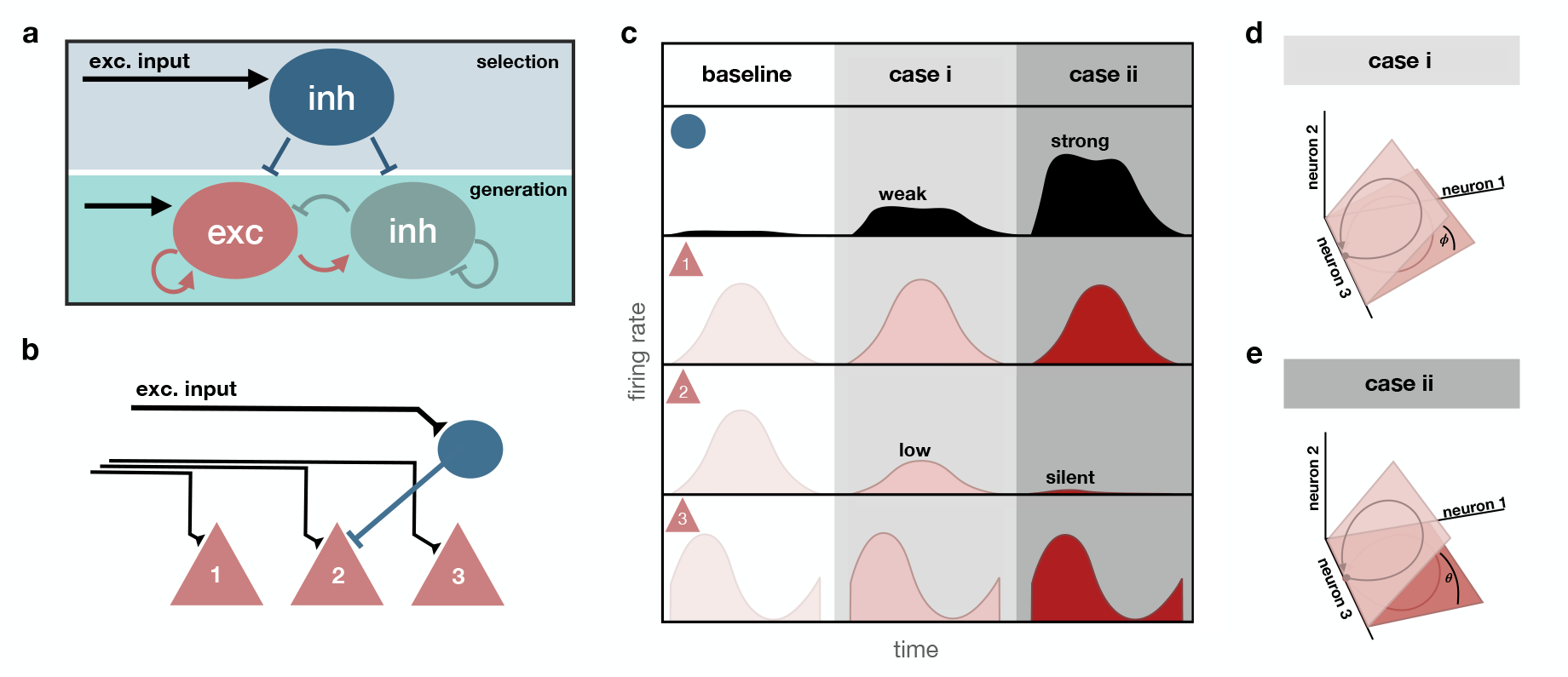
Selective inhibition as a subspace rotation. **(a)** Schematic of a recurrent network of excitatory (red) and inhibitory neurons (green) for sequence generation. A second population of inhibitory neurons (blue) is responsible for sequence selection. Black arrows denote excitatory input. **(b)** Simplified circuit diagram of three excitatory neurons, with neuron 2 receiving selective inhibition. For this example, recurrent interactions are neglected. **(c)** Schematics of firing rates of the selective inhibitory input affecting the excitatory neuron 2, top, and the response of the three excitatory neurons below, as shown in **b**. Three cases for the inhibitory input onto neuron 2 are shown. A baseline condition as well as strong and weak inhibition. Strong inhibitory input silences neuron 2 and weak input lowers the firing rate. **(d**,**e)** Joint activity of the neurons resides on a two dimensional subspace. Colors and numbering correspond to cases in **c. (d)** Inhibition rotates the subspace. **(e)** Silencing neuron 2 rotates the circuit subspace onto the plane spanned by neurons 1 and 2.

### Selectively inhibiting neurons can rotate the neural subspace

Selective inhibition can be understood as a mathematical rotation of the low dimensional subspace on which neural activity unfolds. To illustrate this, we start with a simple example ignoring recurrent connections and consider three excitatory neurons that form part of a sequence (Fig. 1b), one of which receives an inhibitory input. Under baseline conditions, the activity sequence of the three neurons lies on a neural subspace, in this case the activity traces an ellipse on a 2D plane (Fig. 1d, for detailed description see Supplementary Text S1). When weakly inhibited, the neuron’s gain is reduced (case *i*, Fig. 1c), which results in a rotation of the neural subspace (Fig. 1d). If the selective inhibitory input is strong enough, the excitatory neuron is silenced (case *ii*, Fig. 1c) and the activity is projected onto the plane spanned by the remaining active neurons (Fig. 1e), also resulting in a rotation of the neural subspace.

In other words, we see that, in principle, selective inhibition can project neural activity onto a different subspace. A key point is that the old and new subspace can be differentiated by an angle (Fig. 1d,e; see Supplementary Text S1), which can be used by downstream brain regions to decode activity on each subspace as a different neural activity sequence. Next we extend this concept in a recurrent neural network model with which we can test the stability of sequence generation under manipulation of the neural subspace with selective inhibition.

### A network model for inhibition-driven projection onto neural subspaces

To better understand the projection of neural activity by selective inhibition onto different subspaces, we considered a rate-based recurrent network for sequence generation. We use a standard rate-based model (see Methods) with time constant *τ* = 1 and first order numerical integration with step size Δ*t* = 1. The firing rates of the neurons are then given by

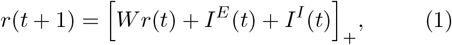

where *r* ∈ ℝ^*N*^ is the vector of firing rates of all neurons in the recurrent network and *W* ∈ ℝ^*N* x*N*^ is the connectivity matrix. Input from the selective inhibitory population is given by *I*^*I*^ ∈ ℝ^*N*^ in addition to the feed-forward excitation by *I*^*E*^ ∈ ℝ^*N*^. The connectivity matrix *W* includes excitation and inhibition responsible for sequence generation on a ring with distance-dependent spatially asymmetric excitation and global inhibition (Fig 2a, see Methods for derivation). This type of connectivity structure gives rise to a *circulant matrix* for the recurrent weight matrix (Fig 2b, see Supplementary Fig. S1), which results in a localized bump of activity forming in the recurrent network. For symmetric distance-dependent excitatory projections following a Gaussian distribution, the activity bump remains stationary. However, if the excitatory projections are made asymmetric, for instance by shifting the center of the Gaussian kernel in one direction, the bump moves around the ring structure, generating a sequence (Fig 2c, Supplementary Fig. S1).

**Figure 2.**
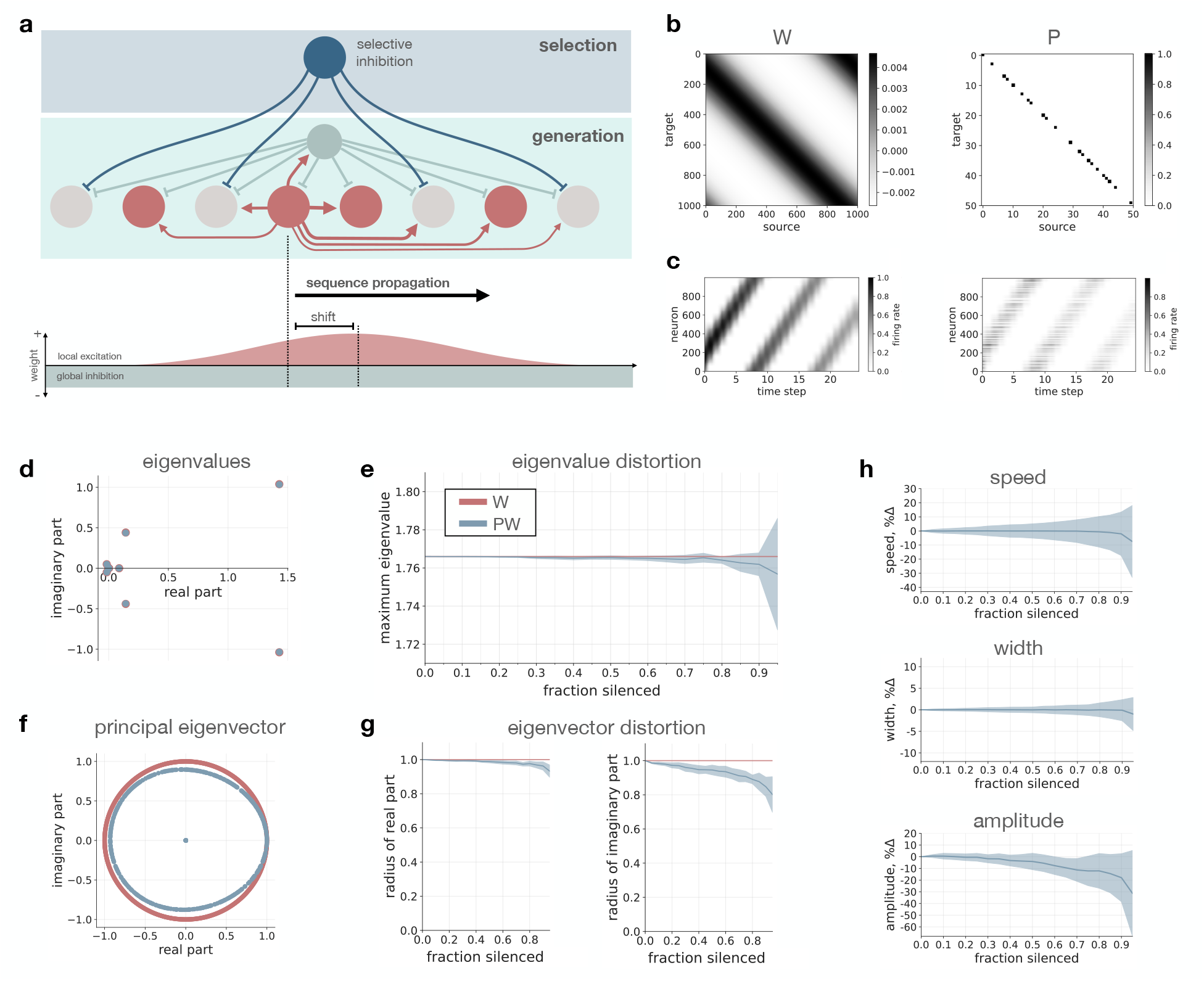
Selective inhibition preserves sequence generation. **(a)** Schematic of network connectivity. Excitatory connections (red) on the ring are local and asymmetric while inhibition (green) is global, leading to sequence propagation. Selective inhibition (blue) targets neurons randomly along the ring. **(b)** Connectivity matrix *W*, left, and an example projection matrix *P*, right. Only 50x50 neurons are shown for visualization. **(c)** Activity bump moving on ring for no projection (*W*), left, and a subspace projection (*PW*), right. Example shows fraction silenced *p*_*inh*_ = 0.6. **(d)** Example eigenvalue spectrum for *W*, red, and *PW*, blue, with fraction silenced *p*_*inh*_ = 0.6. **(e)** Maximum eigenvalue as a function of fraction silenced for *W* and *PW*. **(f)** Principal eigenvector for *W* and *PW* for same example, *p*_*inh*_ = 0.6. **(g)** Radius of real part and imaginary part of normalized principal eigenvector as a function of fraction silenced for *PW*, blue lines. Normalized principal eigenvector for *W* lies on the unit circle, shown for reference in red. **(h)** Percent change in instantaneous speed, width, and amplitude of the activity bump as a function of fraction silenced. In e,g,h line shows mean and shaded region shows standard deviation over 10 different projection matrices.

To this sequence generation network we add a second inhibitory subpopulation, which forms clustered projections onto a subset of neurons in the ring network (Fig. 2a). Neurons receiving inhibition from the second population are uniformly distributed on the ring, meaning the connectivity is uncorrelated with the spatially local connectivity within the sequence generation network.

In order for this selective inhibitory input to generate a subspace projection, we make two assumptions. First, we assume that the inhibitory input remains constant over a time interval *t* ∈ (*t*_1_, *t*_2_), which could range from milliseconds to seconds or longer. This means for a subset of neurons *i* ∈ *S* we have 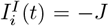, while for the rest *i* ∉ *S* we have 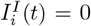. We can rewrite the above equation for the firing rate of neuron *i* as

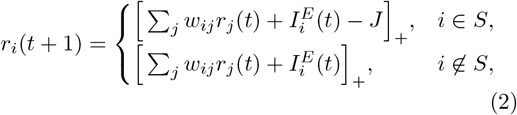

where the sum is over all inputs *j* to neuron *i*.

The second assumption is that the total inhibitory input is strong enough to silence the neurons receiving selective inhibition. For *J* large enough, i.e.

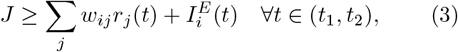

neurons in *S* remain inactive, that is

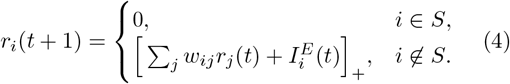

By defining

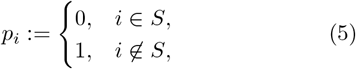

we can rewrite the firing rate update equation as

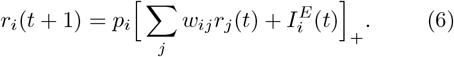

This specifies a projection matrix

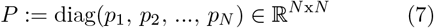

with the *p*_*i*_’s along its diagonal, i.e. zeros in rows of neurons that receive inhibition and ones in rows of neurons that remain active. Thus, Equation 1 becomes

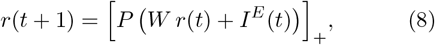

which describes a projection of the circuit activity at each time step onto the subspace spanned by neurons *i* ∉ *S*.

Here we consider the scenario where a transient input triggers bump formation and subsequent sequence progression is intrinsic to the recurrent network. This means that after the bump is initialized, excitatory input ceases (*I*^*E*^ = 0) and the recurrent network locally propagates the sequence. Thus, for Figures 2 and 3 our rate model further simplifies to

**Figure 3.**
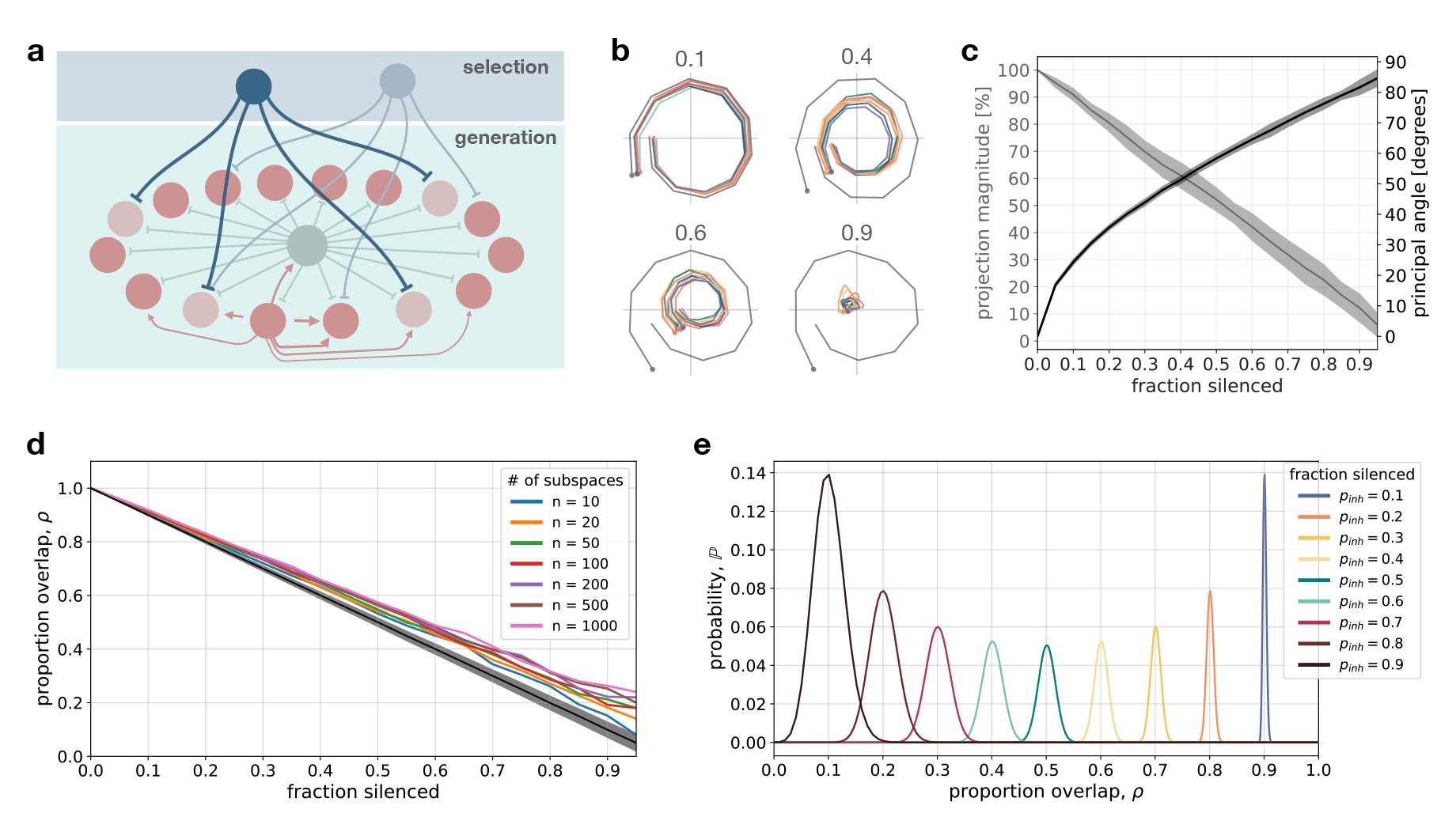
Selective inhibition projects activity onto unique neural subspaces. **(a)** Schematic of ring network for sequence generation and two inhibitory ensembles for sequence selection via projections onto specific subspaces. **(b)** Projections onto PC space for four examples of *p*_*inh*_. Dark grey line shows in-subspace and colored lines show out-of-subspace projections. **(c)** Projection magnitude of out-of-subspace projections and principal angle between subspaces as a function of fraction silenced. Pairwise comparisons for 10 different inhibitory ensembles, i.e. projection matrices *P*. Lines show mean and shaded regions standard deviation over the 45 possible pairs for each fraction silenced. **(d)** Proportion overlap between active neurons in pairs of subspaces as a function of fraction silenced, when *n* = 10 to *n* = 1000 subspaces were stored. Black line shows mean for *n* = 1000 and shaded region standard deviation. Colored lines show maximum overlap over all pairs of subspaces for each number of stored subspaces. **(e)** Probability distribution 𝕡 of two subspaces overlapping by *ρ* for different fractions of silenced neurons.

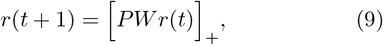

with *r* initialized as a bump of activity at *t* = 0 (see Methods).

Taken together, selective inhibition can be described as a mathematical projection (*P*) acting on the sequence generation network (*Wr*). For this type of subspace manipulation to be useful, it is crucial that the inhibitory projections from the second interneuron population do not significantly impede the network’s ability to generate and propagate sequential activity, which we investigate next.

### Selective inhibition preserves sequence generation

Without selective inhibition, the recurrent model described above robustly generates sequential activity see Supplementary Fig. S1; also see ^26,30,34,76,77^. Sequence generation requires the stable progression between subsequent activity states. In the ring network, this is modeled as the formation of stable bumps of activity and their movement through the network. Bump formation arises due to the distance-dependent excitatory projections and global inhibition, which make the connectivity matrix a *circulant matrix*. The eigenvectors of a circulant matrix are Fourier modes, i.e. complex exponentials, which are spatially periodic ^78,79^. The activity bump arising in the network is determined by the Fourier mode corresponding to the dominant eigenvalue. If the distance-dependent excitatory projections are spatially symmetric, then the connectivity matrix *W* is also symmetric and the eigenvalues are real, which leads to a stationary bump. For asymmetric excitatory projections (i.e. stronger projections in one direction, a shifted Gaussian kernel), positive complex eigenvalues emerge and the bump begins to move through the network (see Supplementary Fig. S1). The magnitude of the imaginary part of the largest eigenvalue is proportional to the “amount of asymmetry”, i.e. shift of the Gaussian kernel, as well as to the speed of bump movement see Supplementary Fig. S1; for similar observations in 2D networks see ^34^.

### Eigendecomposition of weight matrix *W* is preserved under selective inhibition

While the recurrent dynamics without selective inhibition enable sequence generation, we wanted to know whether the projection matrix *P* preserves this property. For sequence generation to be preserved, applying the projection *P* to *W* should not significantly change its eigendecomposition; in particular the dominant eigenvalue and eigenvector should be robust to this projection.

First, analytically we can see that on average the eigenvalues 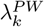 of the matrix *PW* scale linearly as 1 − *p*_*inh*_ times the eigenvalues of *W* with *p*_*inh*_ being the fraction of neurons silenced in the network. In particular,

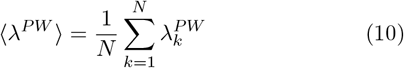

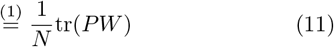

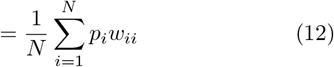

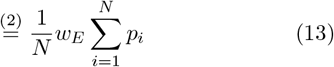

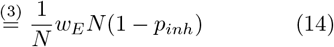

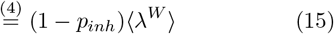

where ⟨ ·⟩ denotes the average and tr(·) denotes the trace of the matrix. The equalities hold because (1) the trace equals the sum of the eigenvalues, (2) the diagonal entries of *W* are all equal to *w*_*ii*_ = *w*_*E*_, (3) the sum of the *p*_*i*_’s is determined by the fraction of silenced neurons *p*_*inh*_, and (4) 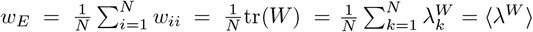.

While this average scaling behavior gives us an intuition, we do not know what will happen to individual eigenvalues, and in particular the dominant eigenvalue-eigenvector pair. To evaluate this numerically, we generated projection matrices *P* to silence a fraction of the neurons from *p*_*inh*_ = 0 up to *p*_*inh*_ = 0.95, in each case for 10 different projection matrices. Due to the silencing, the percentage of zero eigenvalues of *PW* increased linearly from 0% to 95% in a one-to-one correspondence with fraction silenced *p*_*inh*_. The remaining eigenspectrum of *PW* remained highly preserved, with the leading eigenvalues scaling approximately linearly with 1 − *p*_*inh*_ and the structure remaining nearly identical to the eigenspectrum of *W*, even when up to 95% of neurons were silenced (Fig. 2d,e; Supplementary Fig. S3). Furthermore, the principal eigenvector of *PW* was also highly preserved (Fig. 2f,g). The normalized principal eigenvector of *W* lies on the unit circle in the complex plane and applying *P* to *W* only slightly distorted this form (Fig. 2f). We repeated the analysis for a range of asymmetry values and found that the results held, with the largest changes in eigendecomposition for small shifts close to the symmetric case, corresponding to very slow moving or nearly stationary bumps (Supplementary Fig. S2).

To understand why the eigendecomposition is preserved under selective inhibition *P*, it helps to consider that applying the projection *P* is equivalent to setting a subset of rows of *W* to zero. Setting all entries in row *i* to zero corresponds to removing the inputs to neuron *i*, which results in silencing its activity. Setting a row of *W* to zero further implies that the corresponding column of *W* also becomes zero. This follows logically as a column of *W* represents the outputs of the now silenced neuron. We constructed the reduced matrix 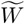 formed by removing the rows and columns of *W* corresponding to the silenced neurons and found that its eigendecomposition is equivalent to that of *PW* (Supplementary Fig. S3). Visually inspecting these reduced matrices makes it clear that due to the uniformly random silencing of neurons, they remain close to circulant (Supplementary Fig. S3), giving further intuition into the preserved eigendecomposition of *PW*.

Thus silencing a uniformly distributed subset of neurons via selective inhibition *P* results in small eigenvalues of the effective weight matrix *PW* becoming zero and the leading eigenvalues scaling linearly, proportional to the fraction of neurons silenced. This means that the structure underlying sequence generation is preserved even when a large fraction of the neurons in the network are silenced via selective inhibition. For the implications of the linear scaling of the eigenspectrum, see the second last paragraph of this section as well as the spiking network simulations in the final results section.

### Sequence progression is preserved under selective inhibition

We now turn our attention away from connectivity to neural activity and ask whether sequence progression is preserved under selective inhibition. To confirm the results comparing the eigendecomposition of *PW* and *W*, we ran simulations to assess the fidelity of sequence progression (bump amplitude, shape, and speed) under subspace projections for fractions silenced from *p*_*inh*_ = 0 to *p*_*inh*_ = 0.95, in each case for 10 different projection matrices, applying the 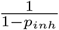 weight scaling as observed above to form *PW*.

We initialized a bump on the ring at *t* = 0 (see Methods) and the firing rates were evolved according to Equation 9. To evaluate sequence progression, for each time step we fit a Gaussian to the activity (see Methods). We computed the amplitude and width (standard deviation) of the fitted Gaussian, as well as its center position, which allowed us to estimate the bump’s speed. In the case without selective inhibition (*p*_*inh*_ = 0), activity bumps have a constant width and speed (Fig. 2h). We adjusted the recurrent weights such that for *p*_*inh*_ = 0 the activity bump decayed in amplitude to half its original height by the end of the simulation.

For each simulation, we computed the instantaneous speed, width, and amplitude of the bump at each time step relative to the values for *p*_*inh*_ = 0. We computed the mean and standard deviation across time steps to quantify how variable the bump’s size and movement is along the ring. On average, the width and speed of the bump remained stable even for a large fraction of silenced neurons, with the variability increasing with fraction silenced. The amplitude was similar for a large range of fraction silenced, decreasing faster on average and becoming more variable for higher fractions of silenced neurons (Fig. 2h). Similar results were found for different shift magnitudes (Supplementary Fig. S4), with sequence progression becoming unstable only for low asymmetry and high fraction silenced. Thus, sequence progression remains stable even when inhibitory inputs densely innervate the sequence generation network, provided the spatial asymmetry is large enough for robust bump movement.

Increased variability in the Gaussian fit to the activity bump along the ring is expected, as despite neurons being silenced based on a uniform distribution on the ring, for each particular projection matrix *P* there is variability in the number of silenced neurons at each spatial location. Variability in neural activity, in this case in the number of active neurons (i.e. population firing rate), is a natural feature of activity in biological brains and not inherently bad in a model, provided that it does not render the network unstable leading to the activity dying out or exploding.

The biological implication of scaling the weights by 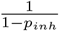 for each fixed fraction silenced deserves some elaboration. There are two possible cases that may arise. If the fraction of silenced neurons *p*_*inh*_ remains the same over time within a network, in other words the density of inhibitory projections is similar across different inhibitory ensembles, then no weight scaling is required and 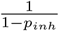 is fixed. However, if projection density differs across inhibitory ensembles then the fraction of silenced neurons would change dynamically. In this case either (i) there must be an additional biological mechanism to dynamically scale the effective weights or (ii) the neuron’s inputs must be saturating, meaning there are sufficiently many synapses for the input to activate the neuron even with fluctuations in the fraction of neurons silenced. In a later section, we will see this case of saturating inputs in a 2D spiking neural network, where large variations in sparsity are possible without requiring weight rescaling.

Taken together, here we have shown that the network dynamics remains stable as evidenced by the eigendecomposition as well as sequence generation and progression remaining preserved even for very large fractions of neurons silenced by selective inhibition.

### Selective inhibition projects sequential activity onto unique neural subspaces

Next we tested our hypothesis that selective inhibition can reorient the neural subspace. For this, we quantified the difference between projections onto different subspaces as a function of the inhibitory sparsity, that is the fraction of neurons silenced. For each fraction silenced, again from *p*_*inh*_ = 0 to *p*_*inh*_ = 0.95, we considered 10 inhibitory ensembles (Fig. 3a), which as described above, equates to 10 different projection matrices. Neurons receiving inhibition from an inhibitory ensemble were again chosen uniformly at random.

As in the example shown in Fig. 1, to quantify differences in subspace orientation we first measured the angle between pairs of subspaces. We ran simulations for each of these cases and, for each activity raster *A* ∈ ℝ^*N* x*T*^, we computed the first three principal components and calculated the first principal angle between pairs of subspaces (Fig. 3c, see Methods). In agreement with our hypothesis, the angle between the subspaces increases with the fraction of neurons silenced, with nearly 90° angles for 95% silencing. Thus, depending on the level of sparsity, clustered inhibition can result in reorientation of the neural subspace from nearly aligned to completely orthogonal, exhibiting a large range of possible behavior.

As a second measure, we considered the projection magnitude of neural activity in each of the neural subspaces (Fig. 3b,c). Each neural activity raster *A* ∈ ℝ^*N* x*T*^ has its own neural subspace in which its projection is maximal. When activity from another activity raster, which lives in a different subspace, is projected onto the subspace belonging to *A*, the relative projection magnitude depends on the alignment of the two subspaces. This measure provides a method for visualization (Fig. 3b) and quantifies the possibility of making an error in decoding, since large magnitude projections in other subspaces makes the trajectories harder to separate. We observed that the relative projection magnitude decreased as the fraction of silenced neurons was increased, ranging from 100%, when no neurons were silenced, to around 5%, when 95% of neurons were silenced. This measure shows that the sparsity of clustered inhibition can determine a full range of behavior, from highly similar trajectories to highly separable trajectories living in orthogonal subspaces. Example subspace projections for different levels of inhibitory sparsity are shown in Fig. 3b.

As a final measure, we considered the amount of overlap in the subpopulation of neurons participating in a sequence across different subspaces. For each inhibitory ensemble delivering selective inhibition, and in turn projecting neural activity onto its corresponding subspace, there is a subset of neurons that are silenced and a subset that remain active. We computed the amount of overlap in recruited neurons between pairs of subspaces as a function of the sparsity of inhibitory projections, i.e. the fraction of neurons silenced. Average proportion overlap decreased linearly with the fraction of neurons silenced by selective inhibition with 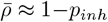 (black line, Fig. 3d). Thus, the denser the innervation from selective inhibitory ensembles (*p*_*inh*_ → 1), the less overlap in recruited subpopulations (*ρ* → 0), paralleling the results for projection magnitude and principal angle.

Next we considered how the overlap depended on the total number of stored subspaces or inhibitory ensembles. Here, the expectation is that as more subspaces are stored, there should be more overlap/alignment between pairs of subspaces. To quantify this, we considered the maximum proportion overlap as a function of the number of stored subspaces. We considered simultaneous storage of 10, 20, 50, 100, 200, 500, and 1000 subspaces. Again, each subspace corresponds to the sequential activity of the neurons in the ring network that remain active under uniformly random silencing of *p*_*inh*_ · *N* neurons by an ensemble of selective inhibition. As expected, we observed that maximum overlap increases with the number of neural subspaces (colored lines, Fig. 3d). However, even for *n* = 1000 subspaces the proportion overlap remains low and still decreases linearly with fraction of neurons silenced. Even with a large number of subspaces, the proportion of neurons shared by any two subspaces remains low when enough neurons are silenced, e.g. maximum overlap is less than 50% when 60% of neurons are silenced by selective inhibition (Fig. 3d). Thus, the large range of behavior in subspace orientation, from aligned to orthogonal, induced by selective inhibitory ensembles is preserved even when many subspaces are stored simultaneously.

To explain this effect analytically, we computed the complementary cumulative distribution function (*ccdf*), which describes the probability that the overlap between two subspaces is more than proportion *ρ* ∈ [0, 1] (see Methods). The *ccdf* is given by

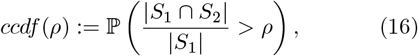

where *S*_1_, *S*_2_ are the set of neurons in subspaces #1 and #2, |*S*_1_ |= |*S*_2_ |:= (1 − *p*_*inh*_) · *N* is the number of active neurons in a subspace (*N*, number of neurons in the network), and |*S*_1_ ∩ *S*_2_ | is the number of neurons that are active in both subspaces (overlap). For subspaces to be distinguishable, the probability of substantially overlapping should be low, though exactly how similar different subspaces can be would depend on brain region, task, and neural coding scheme. This means the *ccdf* should approach zero above some threshold *ρ*_*thr*_

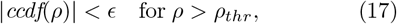

for small positive constant *E* and overlap threshold *ρ*_*thr*_ ∈ [0, 1]. To verify this, we plotted the *ccdf* for different sparsity values from fraction silenced *p*_*inh*_ = 0.1 to *p*_*inh*_ = 0.9 (Fig. 3e). As expected, independent of sparsity, the probability of overlapping goes to zero as *ρ* → 1. With denser inhibitory innervation, meaning a larger fraction of silenced neurons, the *ccdf* goes to zero increasingly fast. For example, when 60% of neurons are silenced by selective inhibition, i.e. *p*_*inh*_ = 0.6, then the chance of two subspaces overlapping more than 50%, i.e. *ρ* = 0.5, is approximately 10^−7^ (note log scale of y-axis, Fig. 3e). A nice feature of the *ccdf* curves is that, for a given *ρ*_*thr*_ and *E*, they clearly show which level of sparsity is required from the selective inhibitory ensembles.

Taken together, by looking at three measures, namely the angle between pairs of subspaces, the relative projection magnitude of trajectories onto other subspaces, and the amount of overlap in recruited subpopulations by different subspaces, we clearly see that a full range of behavior in subspace projections is possible. The important point is that subspace similarity is controlled by the sparsity of projections from the selective inhibitory ensembles onto the neurons that participate in sequence generation. When storing many subspaces, provided the inhibitory inputs cluster on a large enough subset of sequence generation neurons, selective inhibitory ensembles can innervate a random subset of neurons with very low probability of the subspace aligning with another subspace.

### A neural circuit to dynamically select and maintain subspaces

So far we have shown that selective inhibition preserves sequence generation, while at the same time projecting the neural activity onto unique neural subspaces. Next we turn to our hypothesis that inhibitory ensembles with clustered axonal projections onto neurons supporting sequence generation provide a mechanism for *dynamically* selecting and switching between neural subspaces. Since projections onto different subspaces in our model are being performed by the activity of the inhibitory ensembles, we anticipated that switching between subspaces should be possible on the timescale of this neural activity, namely on the order of tens of milliseconds.

To test this hypothesis, we extended the model to include inhibitory ensembles connected reciprocally with the sequence generation network, with sequence selection controlled by a top down signal, possibly from a higher brain region (Fig. 4a). In the extended model, ensembles in the selective inhibition subpopulation inhibit one another, enforcing winner-take-all dynamics, meaning that only one ensemble is active at a time. Global excitatory projections from sequence generation neurons to the selective inhibition subpopulation keeps the winning ensemble active. The underlying connectivity motif is shown in Fig. 4b.

**Figure 4.**
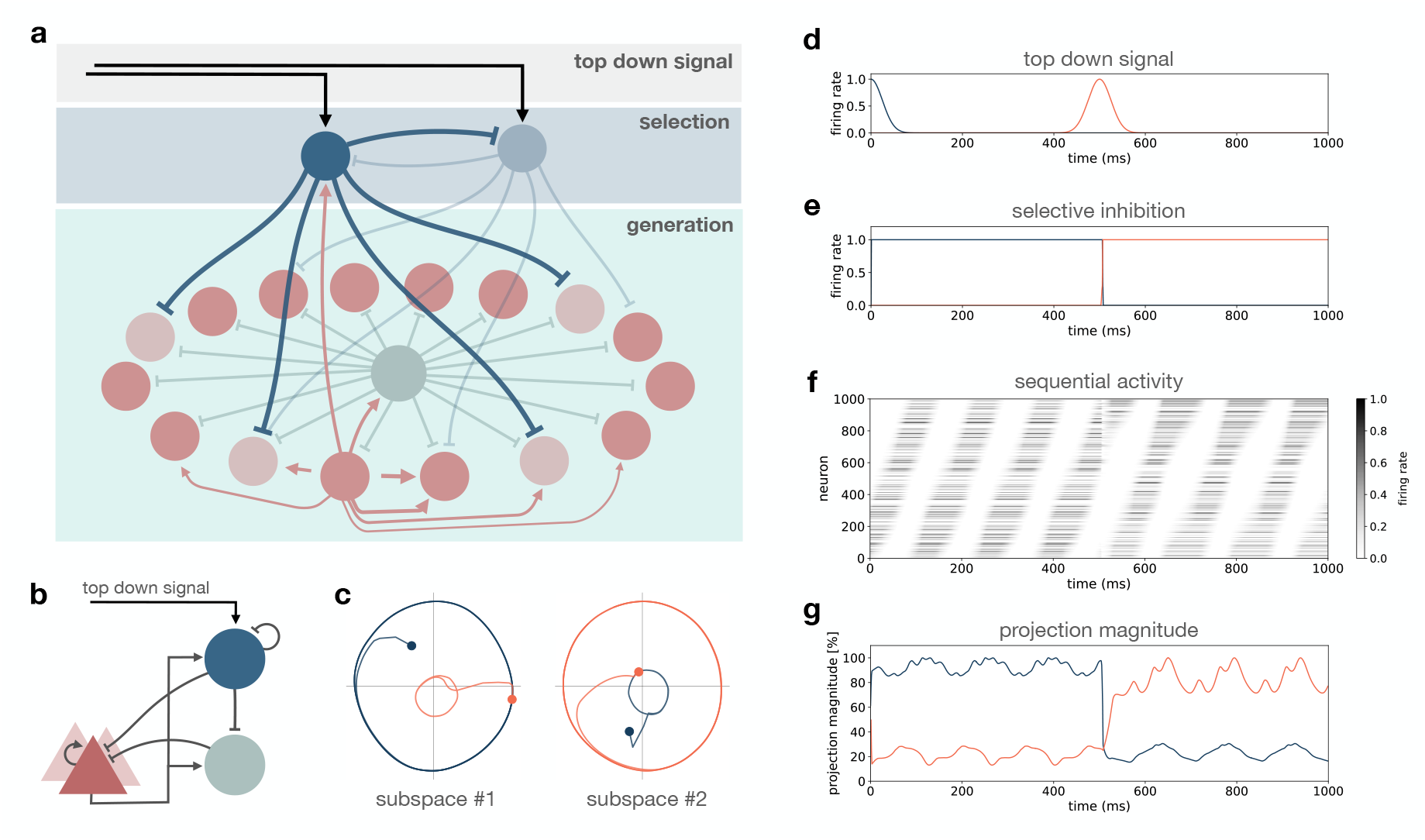
A neural circuit for dynamic subspace selection. **(a)** Schematic of network structure for sequence selection via dynamic rotation of subspaces with selective inhibition. Colors same as previous figures, top down signal, black arrows. Two inhibitory ensembles compete via winner-take-all dynamics to project activity onto their selected subspace. **(b)** Circuit motif underlying connectivity shown in (a). **(c)** Projections of activity from (f) onto neural subspaces. Color code distinguishes periods of time when inhibitory ensemble #1 was active, blue, vs. inhibitory ensemble #2, red. **(d)** Top down signal to selective inhibitory assemblies. **(e)** Activity of selective inhibitory assembles. **(f)** Activity of neurons in the ring network. **(g)** Projection magnitude in subspace #1 and #2, in blue and red, respectively.

We model the firing rate 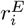 of individual neurons in the sequence generation network as a function of their synaptic currents 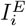 (see Methods for more detail). In particular, we take the following well-established formulation for the rate dynamics ^80^

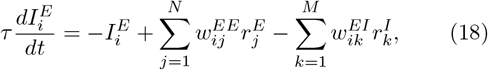

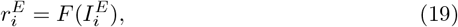

with total synaptic current 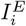 depending on the firing rate of ring network neurons 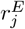 weighted by recurrent weights 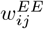 and the firing rate of selective inhibition ensembles 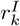 weighted by their inhibitory weights 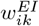.

The synaptic time constant is now set to *τ* = 10ms, extending the results above where *τ* was 1. Recurrent weights 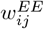 incorporate the local asymmetric excitation and global inhibition required for sequence generation, again forming a circulant connectivity matrix 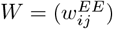 as before. The output firing rate of neuron 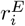 is a function of the total synaptic current 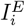, with a piecewise linear activation function *F* keeping the firing rate between 0 and 1. The firing rate of each ensemble of selective inhibitory neurons is modelled similarly as

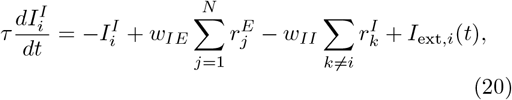

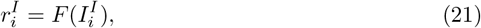

where the total synaptic current 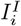 depends on the firing rate of ring network neurons 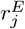 weighted by *w*_*IE*_, the firing rate of the other selective inhibition ensembles 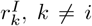, weighted by *w*_*II*_, and top down input *I*_ext,*i*_(*t*). The top down input models excitatory input from an ensemble of neurons as a Gaussian function in time

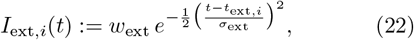

where the firing rate of input ensemble *i* has its peak at *t*_ext,i_ with standard deviation *σ*_ext_, weighted by excitatory weight *w*_ext_.

Within this neural circuit for sequence generation and selection, we then tested the ability to dynamically switch between subspaces. We defined two selective inhibitory ensembles (Fig. 4a) that each project to a random subset of the sequence generation neurons, in this case with *p*_*inh*_ = 0.8, thus leaving 20%, or 200 of the *N* = 1000 neurons active per subspace. Each inhibitory ensemble receives top down input, modeled to resemble the activation of an assembly of input neurons, as described above.

The sequence generation network was initialized with a bump of activity and both inhibitory ensembles set to be quiescent. At *t* = 0 a top down input arrives at the first inhibitory ensemble (Fig. 4d), resulting in it becoming active and suppressing the activity of the second ensemble (Fig. 4e). Activity of the first inhibitory ensemble projects the neural activity sequence onto the according subspace, which is visible in the neural activity (Fig. 4f) as well as in PC space (Fig. 4c). The projection magnitude onto the first subspace is high (blue, Fig. 4g), while the projection magnitude onto the non-selected subspace is low (red, Fig. 4g).

The network remains stable in subspace #1 (blue) and the neural activity sequence unfolds as it should, until a second top down input arrives to the other inhibitory ensemble. This top down input activates the second ensemble, which silences the previously active inhibitory ensemble (Fig. 4e) and projects the activity onto subspace #2 (red, Fig. 4c,f,g). This can be nicely observed in the projections of the activity onto subspace #1 and #2, where the top down input (onset depicted as a red dot, Fig. 4c) forces the neural trajectory off its course in subspace #1, shown in blue, and onto subspace #2, shown in red (Fig. 4c). This sequence selection and dynamic switching is also clearly visible in the projection magnitudes (Fig. 4g), indicating that the switch is successfully executed after about 10-20 ms.

These results show that this neural circuit motif is capable of dynamically selecting and maintaining sequential activity on different neural subspaces, providing a mechanism for fast timescale manipulation of the neural manifold. Notably, this connectivity motif is very stable since the neurons that are active in a given subspace excite the selective inhibition ensemble that in turn silences out-of-subspace neurons. This means that, while the connectivity is reciprocal, it is not a recurrent loop but instead neurons participating in the sequence recruit lateral inhibition of out-of-subspace neurons via the selective inhibition subpopulation.

### Inhibition-driven projection onto neural subspaces in a locally connected spiking neural network

As a last step, we wanted to test whether the circuit motif of spatially heterogeneous clustered inhibition that supported subspace projections in our 1D rate-based ring network would hold in a larger 2D spiking neural network (SNN) for sequence generation. For this purpose, we considered a locally connected SNN, which generates rich spatiotemporal activity patterns via correlated spatial asymmetries in the axonal projections of nearby neurons ^34^, inspired by experimental data ^36–38,65,81–86^. Like in the 1D ring network, neurons make synapses with postsynaptic targets within a local region, with a preference for a particular direction (Fig. 5a). Nearby neurons have a similar preferred direction, and since the network topology is two dimensional, this leads to a smooth 2D landscape of projection asymmetries (Fig. 5a). ^34^ showed that this projection pattern results in feedforward paths through the network that support the generation and propagation of spatiotemporal activity sequences. The model is particularly interesting as its connectivity and activity statistics align with experimental data from cortex ^87^ and the network architecture has been applied to robotic control problems on neuromorphic hardware ^88^.

**Figure 5.**
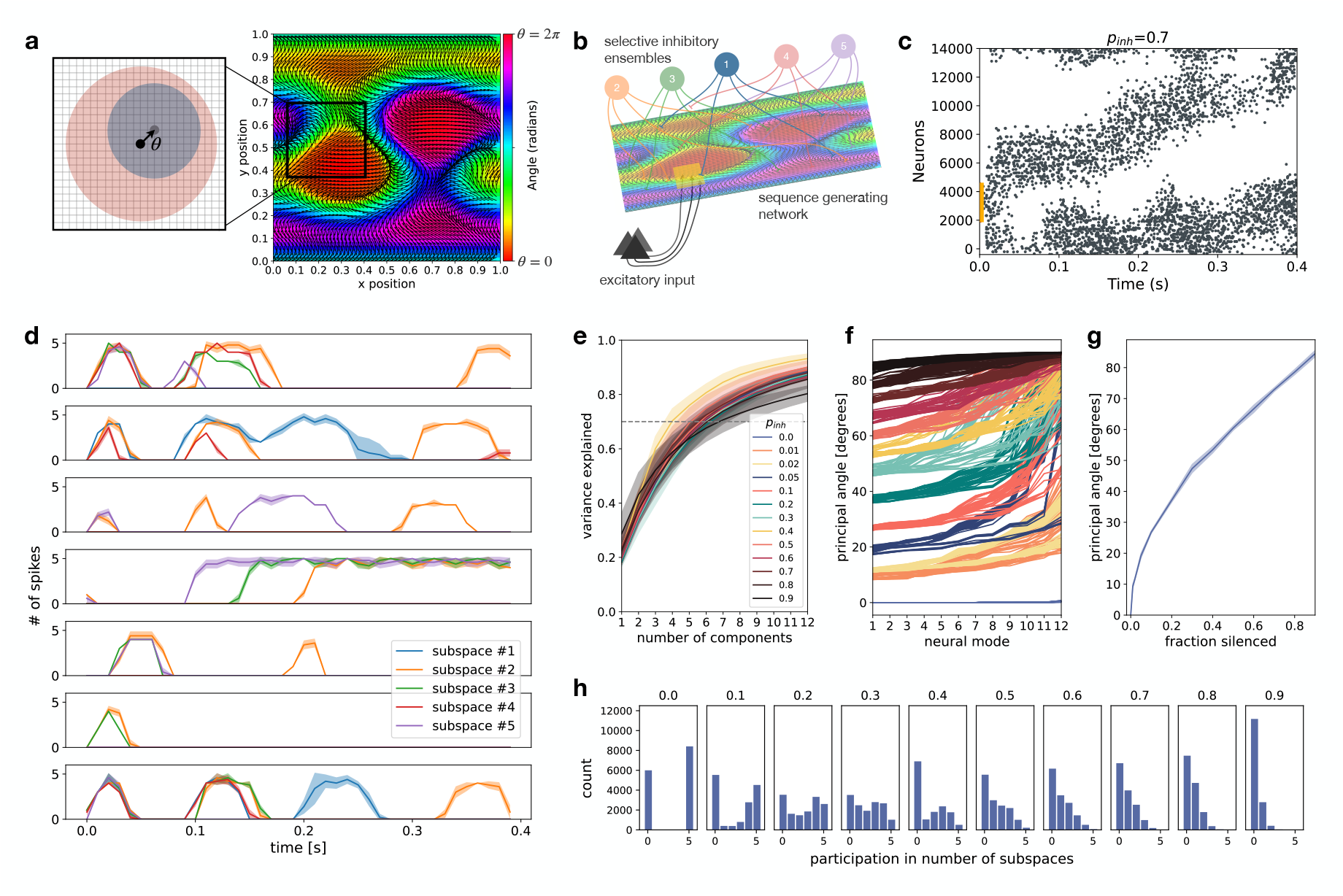
Projection onto neural subspaces in a locally connected spiking neural network for sequence generation. **(a)** Each neuron (left, black point) forms local connections based on a Gaussian kernel (schematic: blue circle, excitatory kernel; red circle, inhibitory kernel). Each excitatory neuron projects in a preferred direction (black arrow), defined by an angle *θ*, implemented as a shift in the center (grey point) of the Gaussian kernel (blue circle). Inhibitory neurons make symmetric projections with a wider Gaussian kernel (red circle). Preferred directions are correlated for nearby neurons (right side, colors denote preferred projection angle *θ* for excitatory neurons on a 2D grid). **(b)** Network structure with selective inhibitory ensembles providing spatially heterogeneous, clustered inhibition silencing a fraction *p*_*inh*_ of the neurons uniformly distributed in the 2D spiking network. Input region (yellow square) receives excitatory input to evoke sequence. Grid depicts the *N*_*E*_ = 14400 excitatory neurons on a 120x120 grid, with landscape of projection asymmetries from (a). (c) Example evoked spatiotemporal activity sequence for fraction silenced *p*_*inh*_ = 0.7. Yellow at *t* = 0 shows input region. Each point represents a spike, every 25th spike is shown. **(d)** Examples of heterogeneous neuron responses in each of the five different contexts. **(e)** Variance explained by the first *k* components for each fraction silenced *p*_*inh*_ ∈ [0, 0.9]. **(f)** Principal angles for the leading 12 principal components (neural modes). Colors denote fraction silenced and each line represents one trial (*n* = 5 trials). **(g)** First principal angle as a function of fraction silenced. **(h)** Histograms demonstrating mixed selectivity for different fraction silenced. Each histogram shows the number of neurons that participated in 0,1,2,3,4 or 5 of the context-dependent spatiotemporal sequences.

To investigate subspace projections via clustered inhibition in the spiking neural network, we conducted a set of simulations in which neural activity sequences were evoked with a transient burst of input to a spatially local region of the network (Fig. 5b,c). We considered a situation in which five different contexts or tasks were stored by five inhibitory ensembles targeting random subsets of *p*_*inh*_ · *N*_*E*_ excitatory and *p*_*inh*_ · *N*_*I*_ inhibitory neurons in the sequence generation network. For each inhibitory ensemble, we simulated five trials in which a sequence was evoked during clustered inhibitory input and white noise background input. We repeated the simulations for a range of inhibitory projection densities, silencing a fraction between *p*_*inh*_ = 0 and *p*_*inh*_ = 0.9 of the neurons for all inhibitory ensembles.

In the spiking neural network model, clustered inhibition projected sequential activity onto unique neural subspaces, and switching between inhibitory ensembles led to rotation of the neural subspace. For each trial, we performed PCA and first measured the proportion of variance explained by the first *k* components (Fig. 5e). This analysis demonstrated that the activity is low dimensional, explained by approximately 4-8 dimensions. We then computed the principal angles between the low dimensional neural subspaces produced by each of the inhibitory ensembles (Fig. 5f). Depending on inhibitory projection density, a wide range of behavior was possible, from close alignment to near orthogonality, in line with observations from monkey motor and prefrontal cortices ^2,11,12^. As in the 1D ring network, the leading principal angle between subspaces increased with the proportion of neurons silenced (Fig. 5g). Thus, the more dense the clustered inhibitory inputs, the larger the angle was between the context-dependent subspaces.

Within individual spatiotemporal sequences, neurons showed rich spatiotemporal behavior (Fig. 5d). Unlike the 1D rate model, neurons were active on different timescales, sometimes more than once within the same sequence, and the behavior of individual neurons (sequential order, timescale, number of times activated) varied across subspaces. Due to the random nature of the clustered inhibition, neurons can be recruited into one or more subspaces, resulting in mixed selectivity across contexts, the extent of which depended on the density of inhibitory projections (Fig. 5h).

Notably, in the spiking network, rescaling the weights as a function of the fraction of silenced neurons (see Equation 15) was not required. When recruited into the sequence, neurons fire a brief burst of action potentials (lasting tens of milliseconds) near or at their peak rate (see Fig. 5d). This means that the input drive is sufficient to push the neuron to saturation. Thus, with sufficient in-degree and sufficiently strong synapses, sequence progression remains stable under balanced silencing of a fraction of excitatory and inhibitory neurons.

While an analytic solution of sequence generation under clustered inhibition in the spiking case is difficult, we can further consider the following reasoning. With connection probability *p* = 0.15 and total excitatory neurons *N*_*E*_ = 14400, average in-degree is *p* · *N*_*E*_ = 2160. Thus, even when 90% of neurons are silenced, on average about 10% or 216 incoming synapses remain viable. Note that neurons may receive multiple synapses from the same presynaptic partner. Based on the excitatory synaptic weights *J*_exc_, a single presynaptic spike increases the postsynaptic voltage by 0.44mV when starting from the resting potential *E*_*L*_ = −70mV. To reach the threshold *V*_t_ = −55mV, the membrane potential must increase by 15mV, and thus requires 15mV*/*0.44mV ≈ 34 simultaneous spikes, neglecting inhibitory inputs and external background input. With a transient burst of input to 20x20 patch of excitatory neurons, as in our simulation, even when 90% of neurons are silenced, 10% of these, or on average 40 neurons, spike due to the input burst. This is sufficient to start sequence propagation. However, as the inhibitory population is recruited and “catches up”, this estimation breaks down and instead we would require an approximation taking the locally correlated asymmetric connectivity and the interaction between excitatory and inhibitory populations into account. While such a general theory for sparse asymmetrically connected 2D spiking networks would be nice, it is difficult in practice. Nonetheless, our simulations show that sequence progression remains stable for a wide range in the sparsity of inhibitory projections.

Taken together, we found that clustered inhibitory inputs silencing uniformly distributed subsets of neurons act to project neural activity onto neural subspaces in a spiking neural network model for sequence generation. Switching between inhibitory ensembles reorients the neural subspace, enabling task- or context-dependent subspaces differentiated by arbitrary angle defined by the inhibitory projection density. Beyond evoked sequential activity, we repeated the experiment for spontaneously generated spatiotemporal activity sequences over a longer timescale of seconds during ongoing background noise and observed similar results (Supplementary Fig. S6).

## Discussion

Given the sequential dynamics of neural activity on task-dependent low dimensional manifolds and the diverse connectivity structures made by the different neuron subtypes within the underlying networks, two important questions arise: (1) how can neural activity manifolds be altered on fast timescales and (2) what kind of computational functions may arise due to different types of inhibitory neurons. In our work we address both of these questions. We have shown that in a recurrently connected network for sequence generation, a second inhibitory neuron subtype providing selective inhibition can project activity onto task- or context-specific neural subspaces. The sparsity of the clustered inhibitory projections controls the angle between subspaces, with a full range of possible behavior from aligned to orthogonal. Importantly, these neural subspaces preserve sequential dynamics, meaning that in each subspace, sequence generation and progression remain intact. Since projections onto subspaces are driven by neural activity of inhibitory cell ensembles, selection and switching can occur flexibly on behaviorally relevant fast timescales. Based on this, we have proposed a neural circuit motif that enables dynamic switching between activity sequences on unique subspaces. Thus, complementing a long list of previous work ^55–61^ we provide additional evidence of how different interneuron subtypes expand the computational repertoire of cortical networks.

### Predictions for connectivity motifs and neural activity patterns

Our model makes specific experimentally verifiable predictions about connectivity motifs and their relationship with neural activity correlations. For example, training an animal on two tasks that require rotation of the neural subspace (as in the task used by ^89^) should result in recruitment of different subsets of interneurons in each task. One subpopulation may be active in all tasks in order to maintain EI balance while another subpopulation/subtype should be selective. The selective inhibitory ensembles should inhibit different but partially overlapping excitatory populations.

Further, our model predicts specific second/third order correlations in II, IE, and EE activity patterns (see Supplementary Fig. S5). For instance, the co-active (co-tuned) inhibitory neurons (correlated in II correlation matrix) should inhibit the same subset of excitatory neurons (correlated in IE correlation matrix) and these excitatory neurons should be distributed throughout the sequence (uncorrelated in EE correlation matrix).

The proposed circuit also predicts an overrepresentation of non-random connectivity motifs. For example, co-active inhibitory neurons should cluster their projections onto a subset of excitatory neurons, which translates to an increase in convergent motifs where multiple inhibitory neurons target the same postsynaptic neuron. There should be more lateral inhibition recruited via the selective inhibition population (i.e. *E*_1_ → *I*_1_ → *E*_2_ but not *E*_1_ ↔ *I*_1_) and more reciprocal connections between excitatory neurons and the inhibitory subpopulation responsible for “EI balance” (i.e. *E*_1_ ↔ *I*_1_).

In our model different groups of inhibitory neurons underlie sequence generation and sequence selection. Identifying these different subpopulations and disrupting their activity should differentially affect the generation of sequential dynamics vs. the selection of a particular sequence/subspace. As such, we predict the separation of sequence generation and sequence selection at different synapses and/or neural subpopulations.

Notably we have assumed that selective inhibition is implemented by ensembles of inhibitory neurons that remain active over a particular time period to perform a subspace projection. It is however conceivable that this second inhibitory population also forms sequences of activity, with each ensemble in the inhibitory sequence providing inhibition to only some of the excitatory neurons forming the activity sequence. This type of scenario may arise if inhibitory projections do not span the entire brain region, but are instead also spatially local.

On that note, this work is motivated by sequential activity in the motor system, hippocampus, parietal cortex, as well as spinal circuits, amongst others. While the model introduced here is not specific to a particular brain region, it could be extended to more closely match a region of interest. Further comparison and integration with experimental results from these regions, like connectivity and activity statistics, and interneuron subtypes, are key steps to verify our findings.

### Fast timescale dynamic switching between activity sequences on unique subspaces

Many situations demand us to dynamically switch between multiple behaviors. This means that dynamically reorienting the neural subspace to enable performance of the appropriate behavior is crucial. So far, computational models have reoriented the intrinsic manifold of a recurrent network using learning ^14,15^. In this setting, reorienting the manifold was not trivial and required either large changes to synaptic weights ^15^ or incremental learning ^14^. However, this type of manipulation of the manifold is slow, and while perhaps feasible for learning new dynamics across many trials or days, cannot explain fast timescale switching between manifolds needed for flexible behavior. Further, when manifold reorientation depends on long-term plasticity at recurrent synapses, the original manifold is lost in the process and dynamic switching between manifolds is not possible. In our model, the projection of neural activity sequences onto neural manifolds is realised by the activity of inhibitory ensembles. Switching between subspaces is therefore fast and dynamic, taking place on the timescale of this activity.

### Clustered inhibition for storage of multiple sequences in one recurrent network

The network model proposed here supports the storage of and selection amongst many neural activity sequences. When clustered inhibitory projections are dense enough, subspace projections have large differences in orientation giving rise to unique subspaces. Critically, we could show that sequence progression is preserved in each of these subspaces. We determined the relationship between inhibitory sparsity and subspace alignment as well as overlap between neuronal subpopulations both in simulation and analytically. Thus for experimental neural recordings of multiple neural subspaces, the model predicts the range of inhibitory sparsity based on the relative orientation of the measured subspaces.

Notably, other computational models based on sequential dynamics in cortex, hippocampus, striatum, and motor circuits have also considered storing multiple sequences. Previous studies have investigated storage of multiple sequences using non-local learning rules ^20,21^, learning at recurrent synapses ^22^, different feedforward excitatory inputs ^24^, gain modulation ^1,90,^, thalamo-cortical loops ^92,93^, multi-chart continuous attractors ^32,94^, or a common sequence that drives different readout networks ^23^. For the models employing thalamo-cortical loops as well as gain modulation via neuromodulatory inputs, the overarching principal is to generate different activity patterns via modulation of the dynamics within a recurrent network by way of a second external mechanism. The inhibitory motif proposed here draws on a similar conceptual argument.

### The order of sequential activation in different sequences

In the 1D ring network, given the local asymmetric connectivity, even when a subset of neurons are silenced by selective inhibition, sequential order of neuron activation remains preserved (i.e. the bump moves around the ring). Preserved co-activation of neurons across conditions (context, decision, behavior) has been observed in a number of experimental settings. Like in our model, in turtle spinal motor circuits two different motor programs give rise to two distinct sequences of neural activity with a highly preserved sequential order of participating neurons see Fig. 5i,j in ^1^. The magnitude of subspace projections across behaviors for these spinal cord sequences also agrees well with our results Fig. 5m in ^1^. In posterior parietal cortex of mice, when making decisions to turn left or right, a subset of neurons are recruited into decision-specific sequences while others are active regardless of the decision with preserved sequential order see Supplementary Fig. 7 in ^3^.

In other studies, partial or complete reshuffling of neuron positions in different sequences has also been observed. In hippocampus, place field arrangements, and hence the order of neuronal activation in hippocampal activity sequences, show a mixture of behavior being partially preserved and partially reshuffling depending on contextual variables ^95–101^. Along these lines, in our 2D spiking neural network, sequential order was no longer preserved across context-dependent subspaces. One contributing factor is that, in the spiking network model, the local asymmetric recurrent connectivity is sparse instead of fully connected. Randomness induced by the sparse local recurrent connections may lead to partial reordering of neural activation while the shift of the kernel still preserves sequential propagation through the network. We also note that, while only neuron-specific inhibition was considered here, we expect that with synapse-specific dendritic inhibition from selective inhibitory ensembles it should be possible to extend the model to shuffle sequence order in a more targeted manner while still preserving sequence generation. With this, a mixture of somatic and dendritic inhibition may control the extent to which neuron order is preserved or shuffled across sequences.

## Conclusion

We considered sequential dynamics in terms of low dimensional manifolds. By leveraging inhibitory diversity, we devised a mechanism by which an inhibitory subpopulation with spatially heterogeneous, clustered synapses dynamically orients the neural manifold, thereby selecting between activity sequences.

## Acknowledgements

We would like to thank Emil Wärnberg for insightful discussions and Michael Fauth for feedback on an early version of the work. We appreciate Leo Hiselius making code related to the spiking neural network implementation publicly available. We thank the Group of Computational Synaptic Physiology for helpful comments. ABL was supported by a Natural Sciences and Engineering Research Council of Canada PGSD-3 scholarship. Nous remercions le Conseil de recherches en sciences naturelles et en génie du Canada (CRSNG) de son soutien. Partial funding from STRATNEURO (to AK) is gratefully acknowledged. We acknowledge support by the Open Access Publication Funds/transformative agreements of the Göttingen University as well as support from the German Research Foundation (Grant number 492788807).

## Data and Materials Availability

All code and data required to produce the results in this paper are available at https://github.com/andrewlehr/subspace projections with selective inhibition.

## Author Contributions

**ABL** developed the idea, conceptualized the study, implemented the simulations, performed the analysis, created visualisations and figures, wrote the original draft, edited/revised the manuscript, and acquired funding. **AK** co-conceptualized the study, provided feedback on the results, edited/revised the manuscript, and acquired funding. **CT** co-conceptualized the study, provided feedback on the results, edited/revised the manuscript, and acquired funding.

