## supplementary_information for "Spatially heterogeneous inhibition projects sequential activity onto unique neural subspaces"

**This PDF file includes:**

Methods
Tables 1 to 4
Supplementary Text
Figs. S1 to S6

### Supplementary Materials

#### 1 Methods

The code to produce all results for this paper will be made publicly available on publication. Parameters for the simulations used to produce the figures in the paper and supplement are listed in Tables 1, 2, 3 and 4.

##### Recurrent network for sequence generation

For the results in Figures 2 and 3, we simulated a recurrently connected network of rate-based neurons. The neurons responsible for sequence generation are organized spatially on a ring. Excitatory neurons form local connections defined by a Gaussian kernel and the inhibitory neurons provide global inhibition. This leads to the following system of equations

$$\tau \frac{dr}{dt} = -r(t) + \Big[W^E r(t) - W^{EI} r^I(t) + I^E(t) + I^I(t)\Big]_+, \tag{S1}$$

$$\tau_I \frac{dr^I}{dt} = -r^I(t) + W^{IE} r(t), \tag{S2}$$

where $r \in \mathbb{R}^N$ is the vector of firing rates of the excitatory neurons in the recurrent network and $r_I \in \mathbb{R}^M$ is the vector of firing rates of the inhibitory neurons providing global inhibition. Here $W^E$ given by the Gaussian kernel $W^E_{ij} = w_E\, e^{\frac{-\Delta x_{ij}{}^2}{2\sigma^2}}$ are the spatially local excitatory recurrent weights (see below, Equation S10), $W^{EI} \in \mathbb{R}^{N\text{x}M}$ and $W^{IE} \in \mathbb{R}^{M\text{x}N}$ are the weights from the global inhibitory population to the excitatory neurons and from excitatory neurons to the global inhibitory population, respectively. $\tau$ and $\tau_I$ are the time constants for the local excitation and global inhibition, respectively. $I^E \in \mathbb{R}^N$ is feedforward excitation and $I^I \in \mathbb{R}^N$ is the input from the second inhibitory population, which forms clustered synapses onto a uniformly distributed subset of neurons in the ring network. The second inhibitory population is described in subsequent sections. Finally, the function $[x]_+ = \max(0, x)$ is a rectifying nonlinearity ensuring that firing rates do not become negative.

For sequence generation in the ring network we incorporated the global inhibition directly into the recurrent weight matrix $W$. This requires the assumption that the timescale of inhibition is much faster than that of excitation. Explicitly, we take $\tau_I \to 0$, which gives

$$r^I(t) = W^{IE} r(t). \tag{S3}$$

With this, the firing rate equations simplify to

$$\tau \frac{dr}{dt} = -r(t) + \Big[(W^E - W^{EI} W^{IE}) r(t) + I^E(t) + I^I(t)\Big]_+ \tag{S4}$$

By letting $W^I := W^{EI} W^{IE} \in \mathbb{R}^{N\text{x}N}$ we can define the recurrent weight matrix $W$, as

$$W := W^E - W^I, \tag{S5}$$

assuming uniform weights $W^{EI}_{ij} = \phi_{EI}$ and $W^{IE}_{ij} = \phi_{IE}$ for all $i, j$, with positive constants $\phi_{EI} \in \mathbb{R}_{>0}$ and $\phi_{IE} \in \mathbb{R}_{>0}$ such that each entry of $W^I$ becomes $W^I_{ij} = M\phi_{EI}\phi_{IE} =: w_I$ with $M$ being the number of projections from the second inhibitory group. As a result, the rate model becomes

$$\tau \frac{dr}{dt} = -r(t) + \Big[W r(t) + I^E(t) + I^I(t)\Big]_+. \tag{S6}$$

Next, substituting the forward finite difference formula for the derivative $\frac{dr}{dt}$ into the differential equation gives

$$\tau \frac{r(t+\Delta t) - r(t)}{\Delta t} = -r(t) + \Big[ W r(t) + I^E(t) + I^I(t) \Big]_+ \tag{S7}$$

and rearranging for $r(t+\Delta t)$ results in the following expression

$$r(t+\Delta t) = r(t) + \frac{\Delta t}{\tau}\Big( - r(t) + \Big[ W r(t) + I^E(t) + I^I(t) \Big]_+ \Big). \tag{S8}$$

By setting $\tau = 1$ and $\Delta t = 1$ we get the following equation for the dynamics of the rate-based neurons

$$r(t+1) = \Big[ W r(t) + I^E(t) + I^I(t) \Big]_+ . \tag{S9}$$

#### Recurrent weight matrix

Neurons are positioned on a ring each with position $x_i \in \{0, 1, 2, 3, ..., N-1\}$. Each neuron receives local excitation and global inhibition. Local excitation follows a Gaussian distribution and the center of the Gaussian kernel is shifted by the shift parameter $s$. When the shift parameter is zero, neurons make symmetric connections in both directions. Non-zero shifts introduce an asymmetry such that each neuron projects preferentially in one direction. The shift parameter is the same for each neuron.

The weight $w_{ij}$ from neuron $j$ to neuron $i$ is given by (see derivation of Eq. S6)

$$w_{ij} := w_E \, e^{\frac{-\Delta x_{ij}^2}{2\sigma^2}} - w_I \tag{S10}$$

where $w_E$ is the maximum excitatory weight, $w_I$ the global inhibitory weight, $\sigma$ defines the width of the Gaussian kernel, and $\Delta x_{ij}$ is the distance between neuron $i$ and the center of the Gaussian kernel for neuron $j$. The kernel defines the spatial distribution of projections from neuron $j$ along the ring to neurons $i$. The center of the kernel for neuron $j$ is given by $c_j := (x_j + s) \bmod N$. The distance between neuron $i$ and the center of the Gaussian kernel for neuron $j$ can be formulated as $\Delta x_{ij} := \min(|x_i - c_j|, N - |x_i - c_j|)$.

The resulting weight matrix is a circulant matrix (see Fig. 2b in main text and Fig. S1 below for visualization). The general form of a circulant matrix is

$$\begin{bmatrix} a_0 & a_{N-1} & \cdots & a_2 & a_1 \\ a_1 & a_0 & a_{N-1} & & a_2 \\ \vdots & a_1 & a_0 & \ddots & \vdots \\ a_{N-2} & & \ddots & \ddots & a_{N-1} \\ a_{N-1} & a_{N-2} & \cdots & a_1 & a_0 \end{bmatrix}. \tag{S11}$$

Each row (respectively column) of the weight matrix is the same, defined by the Gaussian kernel defined above, but shifted by one position and "wrapped" around the circle.

The key parameters of the weight matrix are therefore the shift of the excitatory Gaussian kernel ($s$), the width of the Gaussian kernel ($\sigma$), as well as the peak strength of the local excitation ($w_E$) and the global inhibition strength ($w_I$).

#### Projection matrices

For strong inhibition that silences a subset of neurons $S$, we can write the inhibition as a projection matrix $P$ (for derivation see Main Text). The projection matrices are diagonal, with diagonal entries equal to one for neurons that remain active, and zero for neurons that are silenced. With this, we can rewrite the system of equations for the firing rates as

$$r(t+1) = \Big[P(Wr(t) + I^E(t))\Big]_+. \tag{S12}$$

The projection matrices are determined by one parameter, the proportion of neurons receiving inhibition $p_{inh}$. Neurons receiving inhibition are selected randomly from a uniform distribution. For this, we used `numpy.random.choice` without replacement.

#### Assumption of transient input

For Figures 2 and 3 in the main text we make the assumption that a transient input activates a subset of neurons in the network leading to the generation of a bump of activity. We initialize the firing rates in the network according to

$$r_i(t=0) = p_i\, e^{-\frac{1}{2}\left(\frac{x_i - x_c}{\sigma_0}\right)^2}, \tag{S13}$$

where $r_i(t=0)$ is the initial firing rate of the neuron at position $x_i$, and $x_c$ and $\sigma_0$ define the center and width of the initial activity bump, respectively. For simplicity we set $\sigma_0 = x_c = \sigma$. $p_i$ is the $i^{\text{th}}$ diagonal entry of the projection matrix $P$, which is zero for silenced neurons, $i \in S$, i.e. those receiving selective inhibition, and one otherwise. With this application of the projection matrix $P$ to the initial firing rates, we assume that the selective inhibition acts on the excitatory neurons at time $t = 0$. The initial bump amplitude, when no neurons are silenced, is $r_{max}(t=0) = 1$ and otherwise smaller than one.

#### Assumption of intrinsic sequence progression

We consider the scenario where sequence progression is intrinsic to the recurrent network. This means that after the bump is initialized, throughout sequence progression there is no external excitatory input ($I^E = 0$) and the recurrent network locally propagates the sequence. Thus, for Figures 2 and 3 our rate model further simplifies to

$$r(t+1) = \Big[Wr(t) + I^I(t)\Big]_+ = \Big[PWr(t)\Big]_+, \tag{S14}$$

with $r(t=0)$ defined as above.

#### Scaling the weights

Multiplying the recurrent weight matrix $W$ by the projection matrix $P$ leads to a decrease in the magnitude of the dominant eigenvalue by a factor of $1 - p_{inh}$, where $p_{inh}$ is the fraction of neurons that are silenced. We therefore scaled the recurrent weights by

$$\frac{1}{1 - p_{inh}} \tag{S15}$$

to ensure that the magnitude of the dominant eigenvalue is the same regardless of the sparsity induced by the projection matrix $P$. When more neurons are silenced, each neuron receives fewer excitatory inputs. For this reason, in the simulations testing the fidelity of sequence progression in the 1D ring network, the recurrent weights were scaled up as fraction silenced was increased. Note that this is a uniform scaling to all recurrent weights.

#### Simulations

For Figures 2 and 3 and related Supplementary Figures S1-S4 we performed simulations for 20 different levels of inhibitory sparsity $p_{inh} \in \{0, 0.05, 0.1, ..., 0.95\}$ and 16 different levels of shift magnitude $s \in \{0, 10, 20, ..., 150\}$, leading to 320 different parameter settings. For each of these parameter settings, we performed 10 simulations with a randomly selected subpopulation receiving selective inhibition, i.e. 10 randomly generated projection matrices. Analysis of the eigenvalues, eigenvectors, sequence progression, subspace orientation, and projection magnitude, as described in the next sections, was done on the data produced by these simulations. The parameters for these simulations are described in Tables 1 and 2. For Figure 4 and related Supplementary Figure S5 the simulation parameters are described in Table 3. For the spiking neural network model results presented in Figure 5 and Supplementary Figure S6, the simulation parameters are described in Table 4 and the model is described below in *Spiking neural network model.*

#### Eigendecomposition

We computed the eigenvalues and eigenvectors of matrices $W$, $PW$, and $\widetilde{W}$ using the NumPy linear algebra package in `python`, in particular with `numpy.linalg.eig`. From this we plotted the eigenvalues in the complex plane (see Fig. 2d and Fig. S1g). We computed the magnitude of the complex eigenvalues (see e.g. Fig. S3). From this we determined the largest eigenvalue and its corresponding eigenvector. We also computed the magnitude of the real and imaginary parts of the largest eigenvalue.

#### Eigenvalue distortion

We wanted to quantify the effect of silencing subsets of neurons via selective inhibition on the eigendecomposition of the projected weight matrix $PW$. For each fraction silenced from $p_{inh} = 0$ to 0.95 we computed the largest eigenvalue of $PW$ for 10 different randomly generated projection matrices. The magnitude of the maximum eigenvalue for weight matrix $W$, without silencing i.e. $p_{inh} = 0$, was considered as a reference (Fig. 2e in main text and Fig. S2a). To quantify how the maximal eigenvalue depends on the fraction of neurons silenced, we computed the mean and standard deviation of the largest eigenvalue of $PW$ over the 10 different randomly generated projection matrices. We plotted mean $\pm$ std with the case of no silencing as a reference for the amount of distortion caused by silencing neurons (Fig. 2e). We repeated the analysis for a range of shift magnitudes (Fig. S2a) and also quantified the effect on the real and imaginary parts of the maximum eigenvalue (Fig. S2b,c).

#### Eigenvector distortion

We quantified the effect of silencing neurons on the shape of the principal eigenvector. For a circulant matrix, the eigenvectors are Fourier modes and lay on a circle in the complex plane. For comparison, we normalized the principal eigenvector of $W$ for $p_{inh} = 0$ by dividing by its radius, making the circle a unit circle.

Again we considered 10 different projection matrices for each fraction silenced $p_{inh} = 0$ to 0.95. For each matrix $PW$, we normalized the principal eigenvector by dividing by the maximum absolute value of its components. Then we quantified how different the principal eigenvector was from the expected unit circle, termed eigenvector distortion. We computed the minimum overall radius as the minimum magnitude of the normalized complex eigenvector. We also computed the radius of the real part and imaginary part, which required interpolating the distorted eigenvector on the real and imaginary axes with `scipy.interpolate.interp1d`.

We plotted the mean and standard deviation of the radius of the real and imaginary parts as a function of fraction of neurons silenced (Fig. 2g). We repeated this for different shift magnitudes and show the results in Fig. S2d,e,f.

#### Characterizing sequence progression: speed, amplitude, and width of the bump

To characterize sequence progression we considered the speed at which the activity bump moved around the ring, as well as the width and amplitude of the bump. To do this we fit a Gaussian function to the firing rate data at each time step, taking the periodic boundary conditions into account. We fit Gaussian functions using `scipy.optimize.curve_fit`. The initialization $t = 0$ was removed and estimation of speed, amplitude, and width were performed starting at time step $t = 1$. Specifics of the method for fitting a Gaussian to the data at each time step are described in the provided code.

From the Gaussian fit we get the amplitude, standard deviation, and the center of the Gaussian. Thus, both instantaneous amplitude and width (standard deviation) were used directly from the fit. For speed, we computed the distance between the center of the Gaussian for successive time steps to get the distance travelled per time step. Since time steps were $\Delta t = 1$, instantaneous bump speed is equal to the distance travelled at that time step.

For each simulation, we computed the percent difference in speed, amplitude, and width relative to the case where no neurons were silenced. We then took the average over time steps as well as the standard deviation over time steps to quantify the variability in speed, amplitude, and width induced by silencing neurons. We further averaged over 10 simulations with different projection matrices, repeating this for each fraction silenced $p_{inh}$ (Fig. 2h) as well as for different shift magnitudes (Fig. S4).

#### Principal component analysis

The result of each network simulation is the neural activity of $N$ neurons recorded for $T$ time steps. This activity can be described as a matrix $A \in \mathbb{R}^{N\text{x}T}$, with each column of the matrix containing the activity vector, e.g. firing rates, $r_t \in \mathbb{R}^N$ at time $t$. For each simulation, we performed principal component analysis (PCA) on the standardized firing rate data (transformed to have mean of zero and standard deviation of one). Standardization was done using `sklearn.preprocessing.StandardScaler.fit_transform` and PCA was computed using `sklearn.decomposition.PCA`. Given that around 80% of the variance was explained by the first three principal components (not shown), only these were kept for further analyses.

#### Principal angles

For each simulation, the set of the first $m = 3$ leading principal components describes a three dimensional subspace in the larger $N$ dimensional neural activity space. Thus, for each selective inhibition ensemble, i.e. each projection matrix, there is a corresponding subspace. To compare the orientation of the subspaces generated by different selective inhibition ensembles, we computed the first principal angle between pairs of subspaces.

To compute principal angles we followed the method described by Gallego et al. (2018). Briefly, organizing the leading $m = 3$ principal components of subspace 1 and 2 into $N$ by $m$ matrices $B_1$ and $B_2$ gives us two $m = 3$ dimensional bases describing the two subspaces. Forming the $m$ by $m$ inner product matrix $B_1^T B_2$ and performing singular value decomposition gives

$$B_1^T B_2 = P_1 C P_2^T.$$

The $m$ by $m$ matrix C is a diagonal matrix with the cosines of the principal angles on its diagonal,

$$C = \text{diag}(\cos(\theta_1), \cos(\theta_2), \cos(\theta_3)).$$

The principal angles are ordered from smallest to largest. When the first principal angle is small, the two subspaces are more closely aligned, and when it is large, approaching 90°, the subspaces are orthogonal.

For each fraction silenced, the first principal angle was computed for the 45 pairs of subspaces given by the 10 different selective inhibition ensembles, i.e. projection matrices. We plotted the mean and standard deviation over the 45 subspace pairs for each fraction silenced (see Fig. 3c).

#### Subspace projections

The transpose of an $N$ by $m$ matrix of principal components $B$ multiplied by the $N$ by $T$ firing rate matrix $A$ gives the $m$ by $T$ subspace projection $B^T A$. We took $m = 2$ and computed two dimensional projections (Fig. 3b and 4c). Subspace projections can be either *in-subspace* or *out-of-subspace*. For subspace projection $B_j^T A_i$, where $A_i$ is an activity matrix and $B_j$ is the matrix of principal components corresponding to activity matrix $A_j$, a projection is called *in-subspace* if $i = j$ and *out-of-subspace* if $i \neq j$.

#### Projection magnitude

We measured the magnitude of the 2D trajectory after projecting into a subspace. For each fraction silenced, as described above, we ran 10 simulations each with a different inhibitory ensemble, i.e. projection matrix. We computed *in-subspace* projections for each of the 10 simulations and *out-of-subspace* projections for each of the 45 pairs of different subspaces. We visualized *in-subspace* and *out-of-subspace* projections in Fig. 3b. The absolute projection magnitude was quantified by taking the range of the data in each of the two dimensions ($x_{range} = x_{max} - x_{min}$ and $y_{range} = y_{max} - y_{min}$) and averaging these two values. Then we computed the projection magnitude of out-of-subspace projections as a percentage of the in-subspace projection and plotted this as a function of fraction silenced (Fig. 3c). When no neurons are silenced, by definition, the projection magnitude is 100% and when a subset of neurons are silenced the projection magnitude is less than 100%. For Fig. 4g, where projection magnitude is shown over time, we take the length of the vector in principal component space, $\sqrt{PC1(t)^2 + PC2(t)^2}$, at each time point, normalizing by the maximum projection magnitude for that subspace to convert the values to percentages.

#### Proportion overlap in subpopulations

To measure the overlap in the subsets of neurons participating in different subspaces, we analytically computed the probability that the subpopulation of neurons $S_1$, remaining active for one inhibitory ensemble overlaps with those from a second inhibitory ensemble $S_2$. In particular, we computed the probability distribution over proportion overlap $\rho \in [0, 1]$, given by

$$\mathbb{P}\left(\frac{|S_1 \cap S_2|}{|S_1|} = \rho\right) := \frac{\binom{N}{|S_1|}\binom{|S_1|}{\rho \cdot |S_1|}\binom{N-|S_1|}{|S_1| - \rho \cdot |S_1|}}{\binom{N}{|S_1|}\binom{N}{|S_2|}}, \tag{S16}$$

where $N$ is the number of neurons in the network, $|S_1| = |S_2| := (1 - p_{inh}) \cdot N$ is the number of active neurons in a subspace, and $|S_1 \cap S_2|$ is the number of neurons that are active in both subspaces (overlap). This tells us the probability that the neurons participating in two different subspaces overlap by proportion $\rho$ given the fraction silenced $p_{inh}$.

For two subspaces to be distinguishable, the overlap between them should be small enough. So we are not interested in whether two subspaces overlap by exactly a certain proportion but instead if they overlap by more than some threshold. We therefore computed the complementary cumulative distribution function (ccdf) from the probability distribution. This gives us the desired information as ccdf measures the probability that the subset of neurons participating in two different subspaces overlaps by more than $\rho$. The ccdf is given by

$$ccdf(\rho) := \sum_{\rho_i > \rho} \mathbb{P}\left(\frac{|S_1 \cap S_2|}{|S_1|} = \rho_i\right) = \mathbb{P}\left(\frac{|S_1 \cap S_2|}{|S_1|} > \rho\right), \tag{S17}$$

where the sum is over all proportion overlap $\rho_i > \rho$. For fractions of silenced neurons from $p_{inh} = 0$ to $p_{inh} = 0.9$, we plotted the ccdf as a function of $\rho$ (Fig. 3e).

#### Neural circuit for dynamic subspace selection

To produce the results for Figure 4 in which subspaces are dynamically selected by top down inputs, we extended the model to incorporate winner-take-all dynamics in the selective inhibition population that can be controlled by top down inputs. The following is largely reproduced from the main text with some additional details.

We model the firing rate $r_i^E$ of individual neurons in the sequence generation network as a function of their synaptic currents $I_i^E$. In particular, we take the following well-established formulation for the rate dynamics (Dayan and Abbott, 2005)

$$\tau \frac{dI_i^E}{dt} = -I_i^E + \sum_{j=1}^{N} w_{ij}^{EE} r_j^E - \sum_{k=1}^{M} w_{ik}^{EI} r_k^I, \tag{S18}$$

$$r_i^E = F(I_i^E), \tag{S19}$$

with total synaptic current $I_i^E$ depending on the firing rate of ring network neurons $r_j^E$ weighted by recurrent weights $w_{ij}^{EE}$ and the firing rate of selective inhibition ensembles $r_k^I$ weighted by their inhibitory weights $w_{ik}^{EI}$. Recurrent weights $w_{ij}^{EE}$ incorporate the local asymmetric excitation and global inhibition required for sequence generation, forming a circulant connectivity matrix $W = (w_{ij}^{EE})$ (see Equation $S10$). Explicitly, the recurrent weight $w_{ij}^{EE}$ from neuron $j$ to neuron $i$ on the ring is given by

$$w_{ij}^{EE} := w_E \, e^{\frac{-\Delta x_{ij}^{\,2}}{2\sigma^2}} - w_I, \tag{S20}$$

where $w_E$ is the maximum excitatory weight, $w_I$ the global inhibitory weight, $\sigma$ defines the width of the Gaussian kernel, and $\Delta x_{ij}$ is the distance between neuron $i$ and the center of the Gaussian kernel for neuron $j$.

Instead of a projection matrix, here we explicitly model the selective inhibition ensembles and their projections onto the ring network. As before, we assume strong projections capable of silencing a subset of neurons in the ring network. The size of the subset receiving these projections is determined by the parameter fraction silenced $p_{inh} \in [0, 1)$. Selective inhibition ensemble $k$ thus makes $p_{inh} \cdot N$ connections to neurons in the ring network, each with inhibitory weight $w_{EI}$. That is, the weight $w_{ik}^{EI}$ from selective inhibition ensemble $k$ to neuron $i$ on the ring is

$$w_{ik}^{EI} := \begin{cases} w_{EI}, & \text{if projection exists,} \\ 0, & \text{otherwise.} \end{cases} \tag{S21}$$

The output firing rate of neuron $r_i^E$ is a function of the total synaptic current $I_i^E$, with a piecewise linear activation function $F$ given by

$$F(x) = \begin{cases} 0, & x < 0, \\ x, & 0 \le x \le 1, \\ 1, & x > 1. \end{cases} \tag{S22}$$

The firing rate of each ensemble of selective inhibitory neurons is modelled similarly as

$$\tau \frac{dI_i^I}{dt} = -I_i^I + w_{IE} \sum_{j=1}^{N} r_j^E - w_{II} \sum_{k \neq i} r_k^I + I_{\text{ext},i}(t), \tag{S23}$$

$$r_i^I = F(I_i^I), \tag{S24}$$

where the total synaptic current $I_i^I$ depends on the firing rate of ring network neurons $r_j^E$ weighted by $w_{IE}$, the firing rate of the other selective inhibition ensembles $r_k^I$, $k \neq i$, weighted by $w_{II}$, and top down input $I_{\text{ext},i}(t)$. The top down input models excitatory input from an ensemble of neurons as a Gaussian function in time

$$I_{\text{ext},i}(t) := w_{\text{ext}}\, e^{-\frac{1}{2}\left(\frac{t-t_{\text{ext,i}}}{\sigma_{\text{ext}}}\right)^2}, \tag{S25}$$

where the firing rate of input ensemble $i$ has its peak at $t_{\text{ext},i}$ with standard deviation $\sigma_{\text{ext}}$, weighted by excitatory weight $w_{\text{ext}}$.

#### Spiking neural network model

The spiking neural network model is based on Spreizer et al. (2019), where more details can be found; see also Lehr et al. (2024).

**Neuron and synapse model.** The subthreshold membrane potential $v(t)$ of each neuron in the network is modeled as

$$C_{\text{m}}\frac{dv}{dt} = -g_{\text{L}}\left(v(t) - E_{\text{L}}\right) + I(t) + \mu_{\text{GWN}} + \sigma_{\text{GWN}} \tag{S26}$$

with the leak conductance $g_{\text{L}}$, the membrane capacitance $C_{\text{m}}$, the incoming current $I(t)$, the leak potential $E_{\text{L}}$, and Gaussian white noise with mean $\mu_{\text{GWN}}$ and standard deviation $\sigma_{\text{GWN}}$.

Synaptic currents $I_{\text{syn}}$ elicited by each presynaptic spike are given by

$$I_{\text{syn}}(t) = J_{\text{syn}}\frac{t-(t_{\text{spk}}+t_{\text{delay}})}{\tau_{\text{syn}}}\exp\left(-\frac{t-(t_{\text{spk}}+t_{\text{delay}})}{\tau_{\text{syn}}}\right)H(t-(t_{\text{spk}}+t_{\text{delay}})) \tag{S27}$$

with $J_{\text{syn}}$ the synaptic strength, $t_{\text{spk}}$ denotes the time of a spike in the presynaptic neuron, $\tau_{\text{syn}}$ is the time constant of the synapse, $t_{\text{delay}}$ is the delay between presynaptic spike and post-synaptic response, and $H$ is the unit step function. The incoming synaptic current $I(t)$ of a given neuron at time $t$ is the sum of a transient burst input $I_E$, selective inhibition $I_I$, and the synaptic currents $I_{\text{syn}}$ that the neuron receives from its $N_{\text{syn}}$ recurrent synapses:

$$I(t) = I_E(t) - I_I(t) + \sum_{\text{syn}}^{N_{\text{syn}}} I_{\text{syn}}(t). \tag{S28}$$

Recurrent synaptic weights $J_{syn}$ and connectivity as well as inputs $I_E$ and $I_I$ are described in the next subsections. If $v(t)$ reaches the threshold $V_{\text{t}}$, a spike event occurs at time $t_{\text{spk}}$. Spikes are transmitted from pre- to post-synaptic neuron with delay time $t_{\text{delay}}$. After this event the neuron experiences an absolute refractory period, $\tau_{\text{ref}}$. Following the spike the neuron's potential is reset to the resting potential $V_{\text{r}}$.

**Neuron subpopulations and spatial organisation.** We model a network of excitatory and inhibitory neurons (see EI-network, Spreizer et al., 2019), with a ratio of four excitatory neurons to one inhibitory neuron based on cortical networks (Braitenberg and Schuz, 2013).

When a presynaptic neuron is part of the excitatory subpopulation, $J_{\text{syn}}$, defined in Equation S27, is equal to $J_{\text{exc}} > 0$ and for inhibitory presynaptic neurons $J_{\text{syn}}$ is equal to $J_{\text{inh}}$, defined as:

$$J_{\text{inh}} = -J_{\text{exc}} \cdot g \tag{S29}$$

with $g$ being the strength of the inhibition. Both subpopulations of neurons are homogeneously distributed on a square grid (x, y $\in [0,1]$) with periodic boundaries (i.e. the surface of a toroid).

**Interneuronal connectivity.** Neurons form distance-dependent connections following a Gaussian distribution with a standard deviation $\sigma_{loc,exc}$ for the projections of excitatory neurons and $\sigma_{loc,inh}$ for the projections of inhibitory neurons. For the axonal projections of excitatory neurons, the center of this distribution is shifted away from the position of the neuron. This leads to a preferred axon direction $\phi_i$ for each neuron. The magnitude of the shift is determined by the shift $d$ and is equivalent for all neurons.

The direction of the shift $\phi_i$ is chosen individually for each neuron with Perlin noise, a gradient noise algorithm (Spreizer et al., 2019; Perlin, 1985). With this method, correlated directions are assigned to neurons that are close to each other. The result is an embedded local feedforward structure. Readers are referred to previous publications for further details (Spreizer et al., 2019; Michaelis et al., 2020; Perlin, 1985).

**Clustered inhibition.** A fraction $p_{inh} \in [0, 0.9]$ of the excitatory and inhibitory neurons receive an inhibitory current $I_I(t) = 12500\text{pA}$ which acts to silence their activity. The silenced neurons are uniformly distributed in the network.

**Evoked and spontaneously generated sequences.** For evoked sequences, excitatory neurons in a spatially local region receive a transient burst of excitatory input $I_E(t)$ in addition to the Gaussian white noise background input. The input $I_E(t)$ was applied to a 20x20 region according to Equation S27 with $t_{\text{spk}} = 0$, $t_{\text{delay}} = 0$, $\tau_{\text{syn}} = 5\text{ms}$, and $J_{\text{syn}} = 1.1\frac{(V_t - V_r)}{\Delta V} J_e = 1.1\frac{15\text{mV}}{0.44\text{mV}} 20e\ \text{pA} = 2038.7\text{pA}$. The synaptic current is calculated based on a change in voltage $\Delta V = 0.44\text{mV}$ due to one spike with synaptic weight $J_e = 20e$ pA, thus requiring $\frac{15\text{mV}}{0.44\text{mV}} \approx 34$ simultaneous inputs to increase the voltage by 15mV from $V_r$ to $V_t$, with constant factor 1.1 to ensure $V > V_t$. For spontaneously generated sequences, no transient burst of input was applied, but instead sequences arise spontaneously under Gaussian white noise input.

### 2 Tables

| parameter | value | description |
|---|---|---|
| $N$ | 1000 | number of neurons |
| $T$ | 25 | number of time steps |
| $w_E$ | $\frac{0.0073}{1-p_{inh}}$ | maximum excitatory weight |
| $\sigma$ | 150 | standard deviation of excitatory Gaussian kernel |
| $s$ | 100 | shift magnitude of excitatory Gaussian kernel |
| $w_I$ | $\frac{0.0026}{1-p_{inh}}$ | global inhibitory weight |
| $x_c$ | 150 | initial bump center |
| $\sigma_0$ | 150 | initial bump width (standard deviation) |
| $r_{max}(t=0)$ | 1.0 | initial bump amplitude |
| $p_{inh}$ | {0.0, 0.05, 0.1, ..., 0.95} | fraction silenced |

**Table 1. Parameters used for Figures 2 and 3 and Supplementary Figure 3.** The $N = 1000$ neurons are placed on a ring with grid positions $x_i = \{0, 1, 2, ..., N-1\}$, position 0 and $N-1$ are immediately adjacent to one another. $\sigma$, $s$, $x_c$, and $\sigma_0$ have units in grid points.

| parameter | value | description |
|---|---|---|
| $s$ | {0, 10, 20, ..., 150} | shift magnitudes |

**Table 2. Parameters used for Supplementary Figures 1, 2 and 4.** Note that for the supplementary figures all parameters were as in Table 1 except for the shift magnitude $s$, which was varied between 0 and 150 in steps of 10.

| parameter | value | description |
|---|---|---|
| $N$ | 1000 | number of neurons |
| $T$ | 1000 | number of time steps |
| $\tau$ | 10 | synaptic time constant |
| $w_E$ | 0.05 | maximum excitatory weight between ring network neurons |
| $\sigma$ | 80 | standard deviation of excitatory Gaussian kernel |
| $s$ | 60 | shift magnitude of excitatory Gaussian kernel |
| $w_I$ | 0.005 | global inhibitory weight |
| $w_{IE}$ | 0.02 | weight from ring network neurons to selective inhibition ensembles |
| $p_{inh}$ | 0.8 | fraction of ring network neurons silenced by each selective inhibition ensemble |
| $w_{EI}$ | 10 | weight from selective inhibition ensemble to targeted ring network neurons |
| $w_{II}$ | 25 | weight between selective inhibition ensembles |
| $w_{ext}$ | 25 | weight of top down input signal |
| $\sigma_{ext}$ | 25 | standard deviation of Gaussian activity for top down input signal |
| $t_{ext,1}$ | 0 | peak time of top down input signal from ensemble 1 |
| $t_{ext,2}$ | 500 | peak time of top down input signal from ensemble 2 |
| $x_c$ | 80 | initial bump center |
| $\sigma_0$ | 80 | initial bump width (standard deviation) |
| $r^E_{max}(t=0)$ | 1 | initial bump amplitude, i.e. firing rate |
| $I^E_i(t=0)$ | $r^E_i(t=0)$ | initial synaptic current of ring neuron $i$ is set equal to its initial firing rate |
| $r^I_1(t=0)$ | 0 | initial firing rate of selective inhibition ensemble 1 |
| $r^I_2(t=0)$ | 0 | initial firing rate of selective inhibition ensemble 2 |
| $I_1(t=0)$ | 0 | initial synaptic current of selective inhibition ensemble 1 |
| $I_2(t=0)$ | 0 | initial synaptic current of selective inhibition ensemble 2 |

**Table 3. Parameters used for Figure 4 and Supplementary Figure 5.**

| parameter | value | description |
|---|---|---|
| $N_{\mathrm{E}}$ | 14400 | number of excitatory neurons |
| $N_{\mathrm{I}}$ | 3600 | number of inhibitory neurons |
| $T$ | $0.4\,s, 2\,s$ | duration of simulation |
| $C_{\mathrm{m}}$ | $250\,\mathrm{pF}$ | membrane capacitance |
| $g_{\mathrm{L}}$ | $25\,\mathrm{nS}$ | leak conductance |
| $E_{\mathrm{L}}$ | $-70\,\mathrm{mV}$ | resting potential |
| $V_{\mathrm{t}}$ | $-55\,\mathrm{mV}$ | threshold potential |
| $V_{\mathrm{r}}$ | $-70\,\mathrm{mV}$ | reset potential |
| $\tau_{\mathrm{m}}$ | $\frac{C_{\mathrm{m}}}{g_{\mathrm{L}}}$ | membrane time constant |
| $\mu_{\mathrm{GWN}}$ | $300\,\mathrm{pA}$ | mean of external Gaussian white noise input |
| $\sigma_{\mathrm{GWN}}$ | $25\,\mathrm{pA}, 200\,\mathrm{pA}$ | standard deviation of external Gaussian white noise input |
| g | $6, 8$ | ratio of recurrent inhibition and excitation |
| $\tau_{\mathrm{ref}}$ | $2\,\mathrm{ms}$ | refractory period |
| $\tau_{\mathrm{e}}$ | $5\,\mathrm{ms}$ | time constant for excitatory synapse |
| $\tau_{\mathrm{i}}$ | $5\,\mathrm{ms}$ | time constant for inhibitory synapse |
| $J_{\mathrm{exc}}$ | $20\mathrm{e}\,\mathrm{pA}$ | excitatory synaptic strength |
| $J_{\mathrm{inh}}$ | $-\mathrm{g}\cdot J_{\mathrm{exc}}$ | inhibitory synaptic strength |
| $\sigma_{\mathrm{loc,exc}}$ | $0.1, 0.075$ | gaussian sigma for excitatory projections |
| $\sigma_{\mathrm{loc,inh}}$ | $0.15, 0.1$ | gaussian sigma for inhibitory projections |
| $t_{\mathrm{delay}}$ | $1\,\mathrm{ms}$ | synaptic delay |
| $p_{\mathrm{con_{exc}}}$ | 0.15 | connection probability for excitatory projections |
| $p_{\mathrm{con_{inh}}}$ | 0.15 | connection probability for inhibitory projections |
| $s_{\mathrm{perlin}}$ | $2, 4$ | perlin scale |
| $d$ | 3 | shift of excitatory gaussian connectivity kernel |

**Table 4. Parameters for Figure 5 and Supplementary Figure 6.** Spiking neural network model parameters for the evoked (Fig 5) and spontaneous (Fig S6) sequences. Where parameters differ for evoked/spotaneous they are listed in order.

### 3 Supplementary Text

#### S1 Selective inhibition can rotate and scale the activity subspace

We want to understand how selective inhibition onto a network of neurons can shape the low dimensional neural manifold. To get some intuition for how this might work, let us start with just a few neurons, such that we can easily visualise what is happening, and then work our way up to larger networks. The main intuition we hope to develop here is that changes in firing rate amplitude can scale and rotate the resulting activity manifold while still allowing activity sequences. Note that here we focus on linear subspaces.

We start by considering a network of excitatory and inhibitory neurons (see Fig. 1a in main text). Excitatory projections from an upstream region exert feedforward excitation onto the excitatory neurons and feedforward inhibition via one type of inhibitory neurons (blue circles). Inspired by "quasi-periodic" patterns in motor cortex, we suppose each excitatory neuron's firing rate can be approximated by a sinusoidal function

$$r_i(t) := A_i(1 + \sin(2\pi f_i t + \varphi_i)), \tag{S30}$$

where $A_i$ is the amplitude, $f_i$ is the frequency, and $\varphi_i$ the phase. Note that the particular response pattern is not important and could be some arbitrary function.

In the case of just two neurons with similar responses (e.g. neurons 1 and 2 in Fig. 1c, main text, with $f_1 = f_2$ and $\varphi_1 = \varphi_2 = -\frac{\pi}{2}$), the firing rates of the two neurons are constrained to a line. Adding a third neuron with a different response phase (e.g. $\varphi_3 = 0$) parameterizes an elliptical trajectory on a 2D plane. Some insight can be gained by writing down the firing rate equations for the three excitatory neurons in matrix form. In particular, we construct a diagonal matrix of the amplitudes, and write

$$\begin{bmatrix} r_1(t) \\ r_2(t) \\ r_3(t) \end{bmatrix} := \begin{bmatrix} A_1 & 0 & 0 \\ 0 & A_2 & 0 \\ 0 & 0 & A_3 \end{bmatrix} \begin{bmatrix} 1 + \sin(2\pi f_1 t + \varphi_1) \\ 1 + \sin(2\pi f_2 t + \varphi_2) \\ 1 + \sin(2\pi f_3 t + \varphi_3) \end{bmatrix}, \tag{S31}$$

which for the neurons in Fig. 1 is $A_1 = A_2 = A_3$, $f_1 = f_2 = f_3$, and $\varphi_1 = \varphi_2 = -\frac{\pi}{2}$, $\varphi_3 = 0$. The diagonal matrix of amplitudes is a scaling matrix, which by scaling each of the dimensions (i.e. neurons' firing rate) controls the shape of the trajectory. This, in turn, controls the orientation of the activity manifold. In the linear case, a change in orientation of the activity manifold due to a change in one or more neuron's firing rate can be quantified by the angle between the subspaces. This measure of the "difference between subspaces" is important when considering how a downstream region might read out patterns of activity, with larger angles implying larger differences between patterns, rendering them more distinguishable.

**Angle between subspaces.** To compute the angle between the subspaces analytically for the three dimensional example, we first eliminate time in the parametric equations. We introduce variables $x_1$, $x_2$, and $x_3$ for the three dimensions (i.e. neurons). In this case the 2D plane is described by the equation

$$A_2 x_1 - A_1 x_2 = 0. \tag{S32}$$

To denote a particular plane $i$, we add a superscript to the amplitudes, e.g. neuron $k$ in plane $i$ is $A_k^i$. The angle between two planes is given by

$$\Theta_{ij} := \arccos \frac{|n_i \cdot n_j|}{||n_i||\,||n_j||}, \tag{S33}$$

where $n_i$ and $n_j$ are the corresponding normal vectors to planes $i$ and $j$. These can be read from the coefficients of the equation of the plane (Equation S32), which gives

$$n_i := \begin{bmatrix} A_2^i \\ -A_1^i \\ 0 \end{bmatrix}, \quad n_j := \begin{bmatrix} A_2^j \\ -A_1^j \\ 0 \end{bmatrix}. \tag{S34}$$

With this we can connect changes in firing rates to changes in the orientation of the activity subspace.

**Effect of inhibition on subspace orentiation.** Let us now consider feedforward inhibition acting on the excitatory neurons. For simplicity, at baseline we assume all amplitudes are equal to one ($A_1 = A_2 = A_3 = 1$). Under inhibition, we suppose neuron 1 has amplitude $0 \leq \alpha \leq 1$ and neuron 2 amplitude $0 \leq \beta \leq 1$. The stronger the inhibition the lower are the values of $\alpha$ and $\beta$. Note that changes to the amplitude of neuron 3 do not change the angle of the plane (cf. Equation S33 and S34), but instead just change the trajectory on the plane. With this, the normal vectors to the planes from Equation S34 become

$$n_1 := \begin{bmatrix} 1 \\ -1 \\ 0 \end{bmatrix}, \quad n_2 := \begin{bmatrix} \alpha \\ -\beta \\ 0 \end{bmatrix}. \tag{S35}$$

Under these conditions we can look at the effect of different levels of selective inhibition on the activity subspace defined by the three excitatory neurons. We distinguish between the following cases:

***Case I*: Scaling.** If all neurons receive the same amount of inhibition ($\alpha = \beta$), all dimensions are scaled to the same degree. The orientation of the subspace does not change, just the trajectory is scaled. A linear subspace is invariant to this type of change and we can see this by computing the angle between the subspaces

$$\Theta = \arccos \frac{2\alpha}{\sqrt{2}\sqrt{2\alpha^2}} = \arccos 1 = 0^\circ.$$

With an angle of $0^\circ$ we see that the two subspaces are perfectly aligned.

***Case II*: Rotation.** A second possibility is that neurons receive different levels of inhibition, but they are not silenced ($0 < \alpha \leq 1$, $0 < \beta \leq 1$, $\alpha \neq \beta$). In this case, depending on the relative level of inhibition to each of the neurons (Fig. 1c, case *ii*, in main text), and hence the relative amplitude of their responses, the activity subspace is rotated (Fig. 1e). For $\alpha = 1$ and $\beta < 1$ the angle of rotation can be computed again with Equation S33 resulting in $\Theta < 45^\circ$ since

$$\Theta = \arccos \frac{1+\beta}{\sqrt{2}\sqrt{1+\beta^2}} < \arccos \frac{1}{\sqrt{2}} = 45^\circ.$$

***Case III*:** If inhibition is strong enough to completely silence neuron 2 ($\alpha = 1$, $\beta = 0$), the result is a rotation of the subspace into the plane spanned by the remaining active neurons (see Fig. 1d in main text). This can be understood by looking at the matrix of the amplitudes in Equation S31 and considering $A_2 = 0$, meaning only the other two dimensions (i.e. neurons) remain. Noteworthy is that this matrix is an orthogonal projection matrix (i.e. $P^2 = P = P^T$) exactly when its diagonal entries are exclusively ones and zeros, i.e. there is no partial inhibition, only complete silencing. Plugging the corresponding normal vectors into the angle formula (Equation S33) gives that the angle between the subspaces for the baseline vs. the projected subspace is

$$\Theta = \arccos \frac{1}{\sqrt{2}} = 45^\circ.$$

**Summary.** So we have been able to understand the concept for two and three dimensions and see that inhibiting neurons can rotate the subspace on which the activity lives. We want to extend this idea to arbitrary network sizes. In particular, what is important to note is that, with more neurons and recurrent interactions betweent them, dimensions of the activity manifold are no longer single neurons, but instead neural modes. These are groups of neurons that tend to be active together, each forming one dimension of the activity manifold.

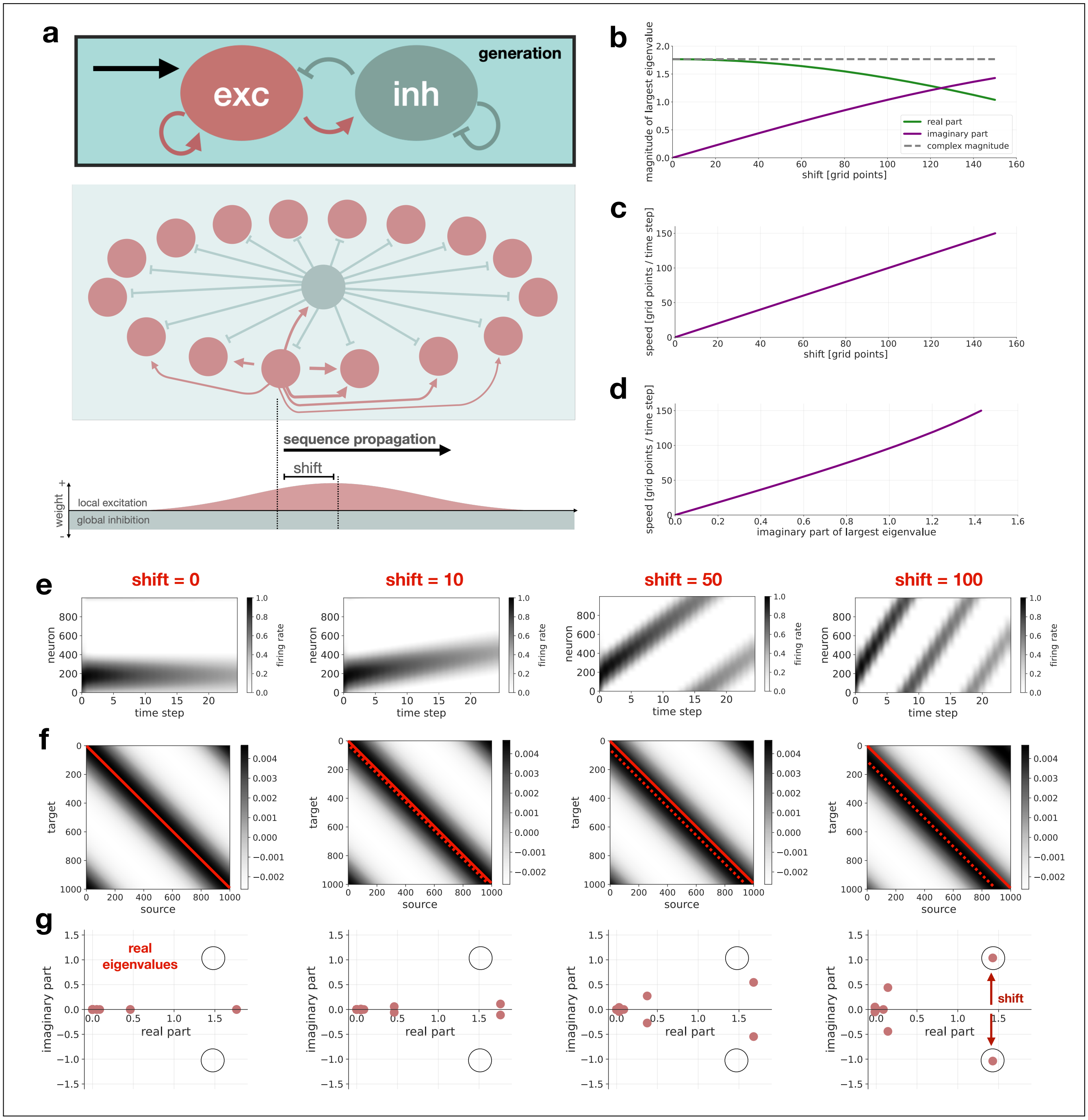


**Supplementary Figure S1. Sequence generation in locally connected ring network.** *Caption on the following page.*

**Supplementary Figure S1. Sequence generation in locally connected ring network.** **(a)** Schematic of network structure with local excitation and global inhibition. When a shift in the Gaussian connectivity kernel is introduced, the sequence propagates through the network. Excitatory connections are shown for one example neuron. **(b)** Magnitude of largest eigenvalue $\lambda_{max}$ of recurrent weight matrix $W$ is shown, as well as the magnitude of the real and imaginary parts, as a function of the shift magnitude. Grid points refer to the positions of the neurons on the grid $x_i = \{0, 1, 2, ..., N-1\}$. The magnitude of the imaginary part increases with shift magnitude. **(c)** Sequence speed and shift magnitude have a one-to-one linear relationship. **(d)** Sequence speed becomes faster as the magnitude of the imaginary part of the largest eigenvalue increases. **(e-g)** The activity, recurrent weight matrix, and eigenspectrum are shown for four different shift values. Activity rasters show that sequence speed increases with increasing shift. The amount of shift is visible in the weight matrices (dashed red line shows shifted center, solid red line is unshifted center as reference). Accordingly the imaginary part of the largest complex eigenvalue pair increases with shift magnitude (black circles show position of largest eigenvalues for shift=100 for reference). When there is no shift, the eigenvalues are real and the bump does not propagate.

---

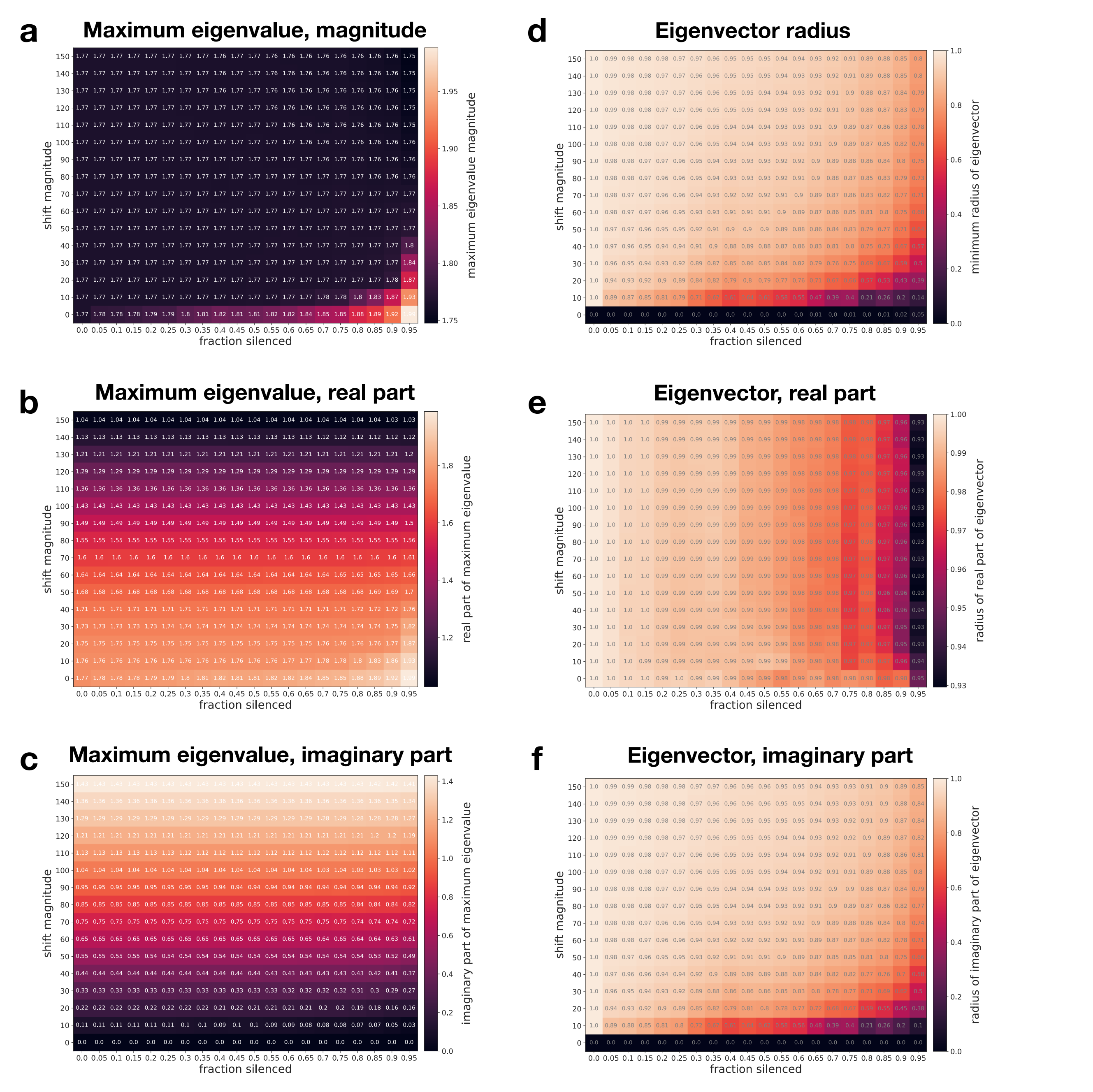


**Supplementary Figure S2. Eigendecomposition of PW as a function of connection asymmetry and fraction silenced** (related to Fig. 2defg in main text). The eigendecomposition of $PW$ was computed for weight matrices $W$ with different levels of connection asymmetry, i.e. shift magnitude from 0 to 150, and projection matrices $P$ with fraction silenced from 0 to 0.95. Note that colorbars in each plot are different. Mean values over 10 different randomly generated projection matrices are shown. The plots demonstrate that the eigendecomposition is robust to silencing of neurons across a large range of connection asymmetry, i.e. shift magnitudes. The following is shown: **(a)** Magnitude of the maximum eigenvalue as well as the magnitude of the **(b)** real and **(c)** imaginary parts. **(d)** Minimum radius of the eigenvector corresponding to the maximum eigenvalue (for no silencing of neurons eigenvector lays on unit circle). Radius of the **(e)** real and **(f)** imaginary parts are shown separately as well.

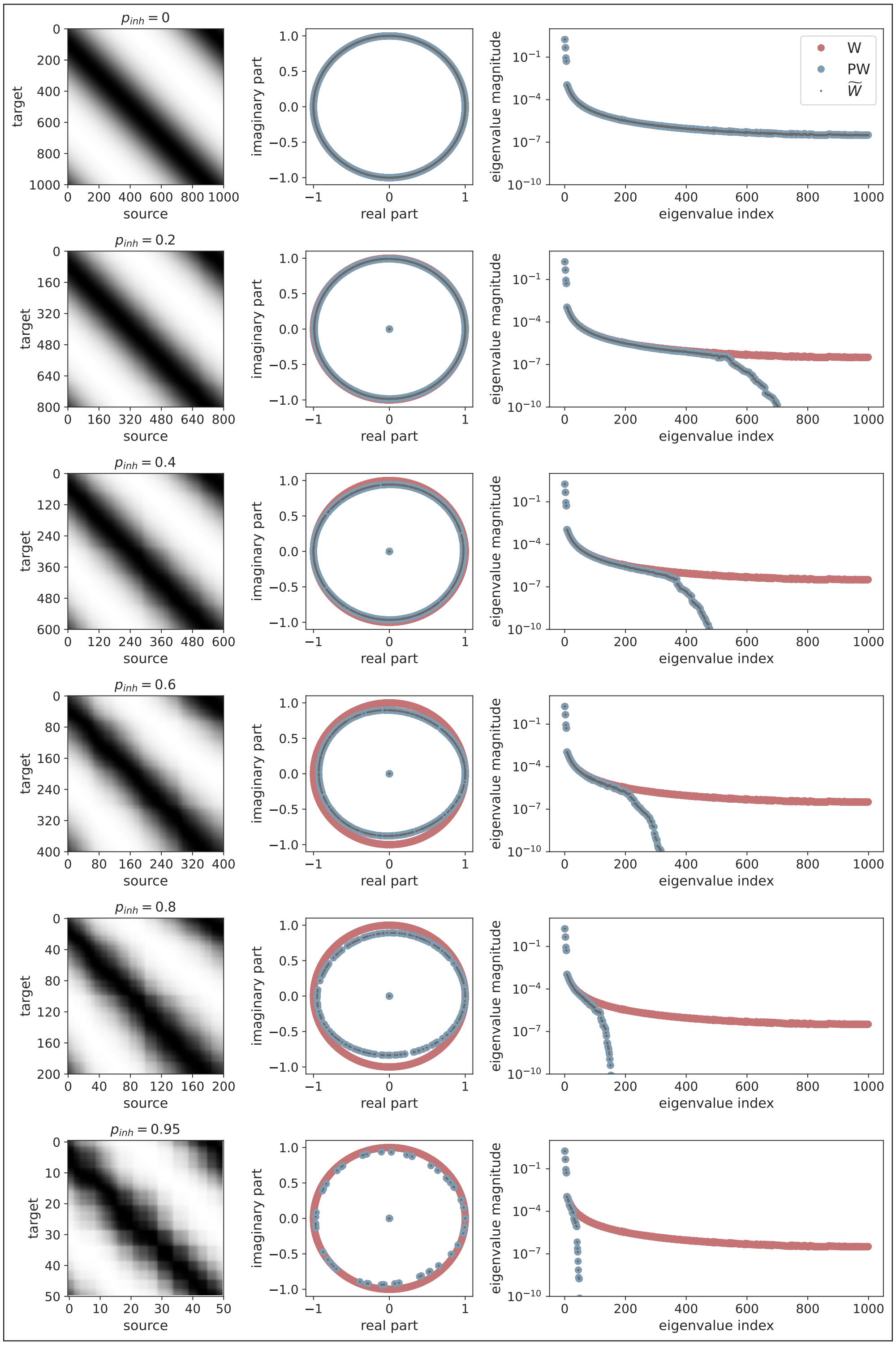


**Supplementary Figure S3. Eigendecomposition of $\widetilde{\mathbf{W}}$ and PW.** The left column shows the reduced matrix $\widetilde{W}$ for different fraction silenced, $p_{inh} = 0$ to 0.95. Note that the matrices continue to look similar to circulant matrices but noisy. The corresponding eigenvectors and eigenvalues are shown in the middle and right columns, respectively. We see that $\widetilde{W}$ and $PW$ are the same. Eigendecomposition of $W$ is shown for reference.

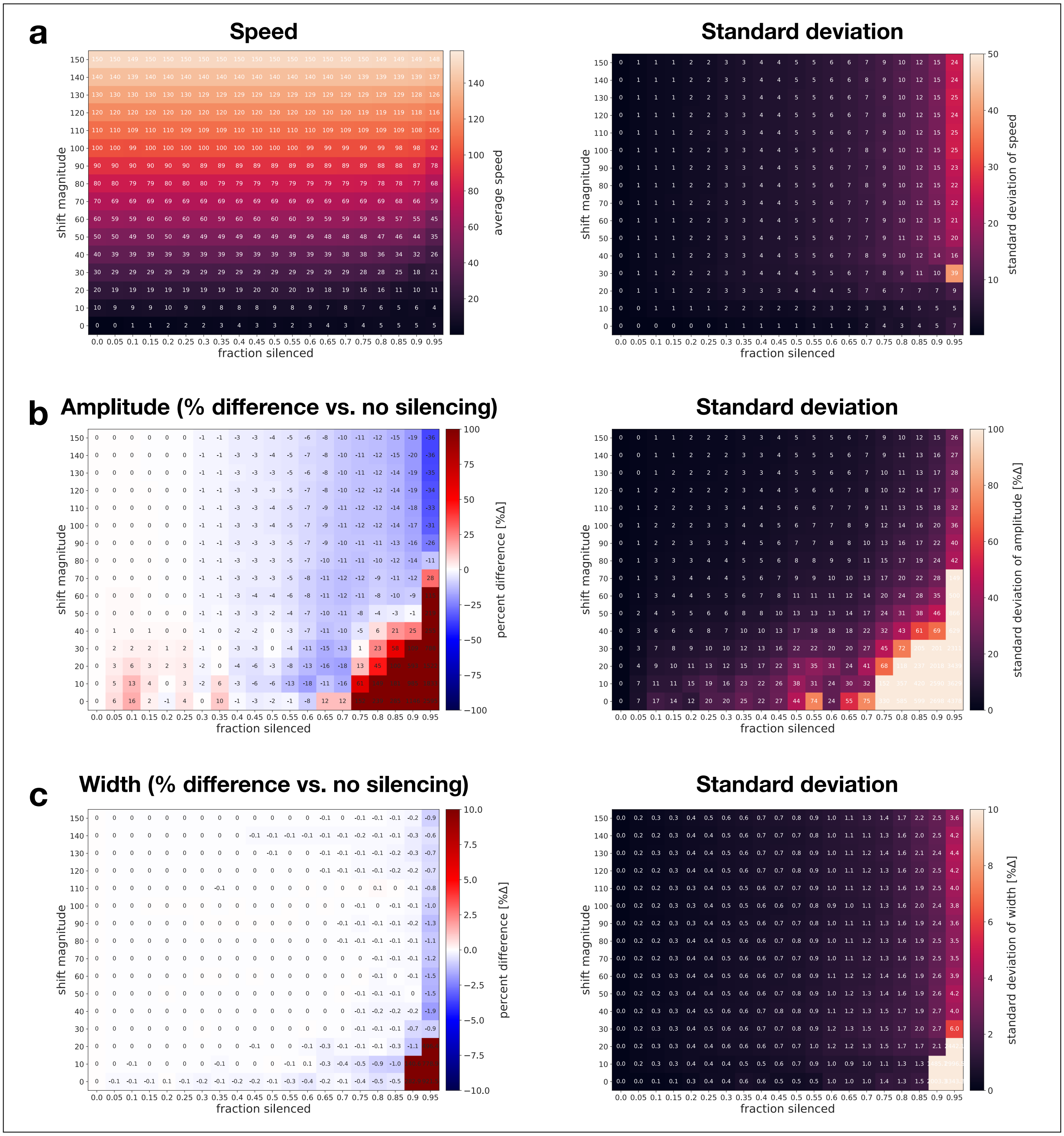


**Supplementary Figure S4. Activity statistics of bump for different shift magnitudes as a function of fraction silenced** (related to Fig. 2h in main text). *Caption on the following page.*

**Supplementary Figure S4. Activity statistics of bump for different shift magnitudes as a function of fraction silenced** (related to Fig. 2h in main text). Results for different shift magnitudes and fraction silenced are shown. For each grid point, 10 simulations were performed each with a different randomly generated projection matrix. An average was computed over time for each simulation and the 10 simulations were then averaged, left column. The right column shows the corresponding standard deviation. Note that the values are stable over a large range of simulations and instabilities arise only for low shift magnitudes (slow moving bump) when a large number of neurons are silenced. **(a)** The bump speed is directly proportional to shift magnitude when no neurons are silenced and is only very marginally affected by silencing neurons. Note in the main text we show percent differences to the baseline case of no silencing, but here we show absolute speeds as these are related to the effect of shift magnitude demonstrated on bump speed shown in Supplementary Fig. S1 above. For **(b)** amplitude and **(c)** width of the bump, as in the main text the percent difference between a given simulation and baseline case with no silenced neurons is shown.

---

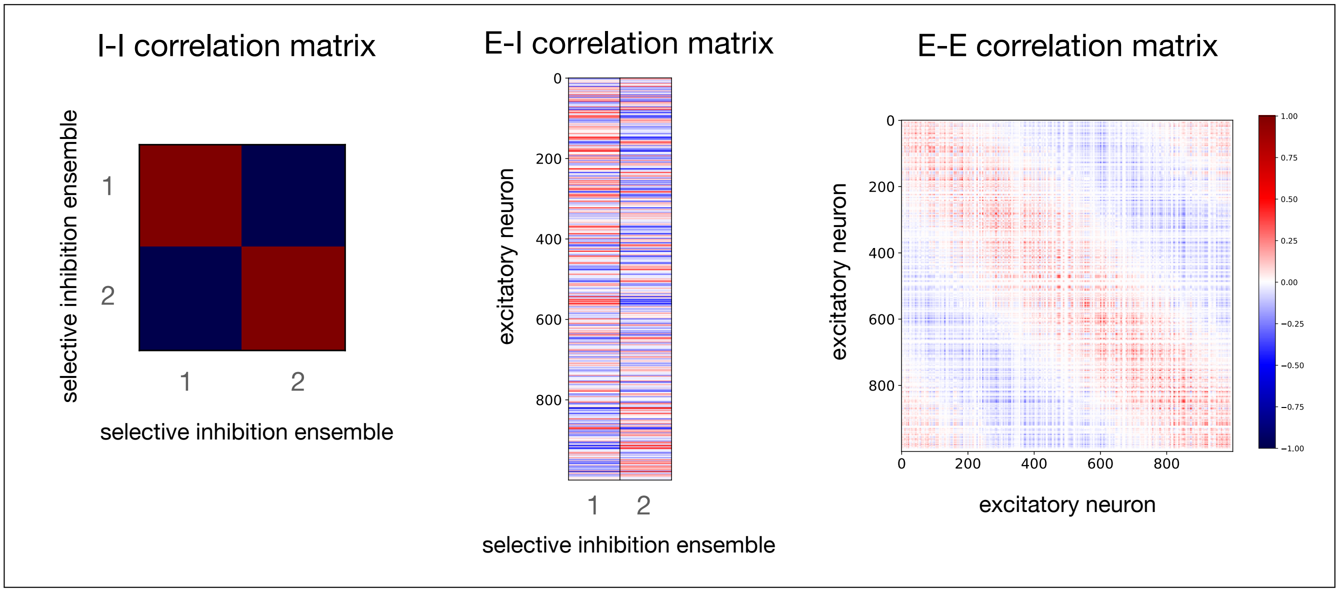


**Supplementary Figure S5. Correlation matrices for II, EI, and EE neurons.** Our model makes predictions about the correlation structure within the subpopulation of inhibitory neurons providing selective inhibition (II), the excitatory neurons responsible for sequence generation (EE), and between the two populations (EI). Shown are II, EI, and EE correlation matrices computed from the data presented in Fig. 4 in the main text, where two selective inhibition ensembles generate different subspace projections. Blue shows negative correlation and red positive correlation, scale from $-1$ to $+1$, with white representing zero correlation. For II interactions, the model predicts that the correlation between neurons in the same selective inhibition ensemble should be strongly positive while neurons in different selective inhibition ensembles should be anti-correlated. Note that here we assume the ensembles are orthogonal. For EI interactions, inhibitory neurons in the same ensemble should inhibit the same subset of excitatory neurons responsible for sequence generation, due to their clustered projections. The model therefore predicts that co-active (co-tuned) inhibitory neurons should have similar structure in their correlations with the excitatory neurons. When the EI correlation matrix is sorted by selective inhibition ensemble there should be horizontal bands corresponding to the neurons targeted by clustered projections (anti-correlated, blue) and the neurons that do not receive inhibition from that ensemble (correlated, red). For the EE correlation matrix, when sorted by the temporal order of neurons in the sequence, a diagonal band is visible, corresponding to co-active neurons within the sequence. The neurons silenced by the selective inhibition ensembles are scattered randomly throughout the sequence, here seen as white vertical and horizontal bands.

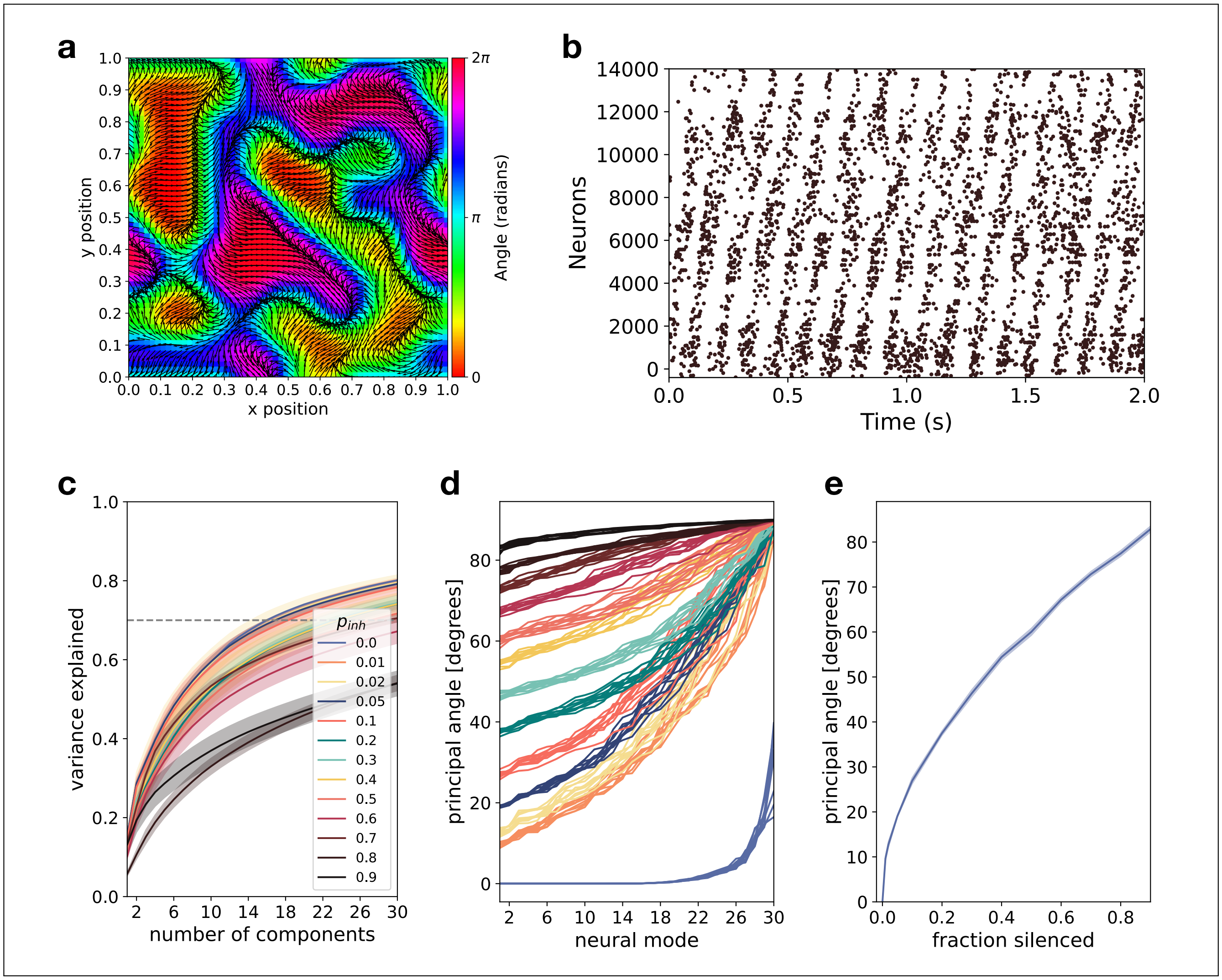


**Supplementary Figure S6. Projection onto neural subspaces in a spiking neural network during spontaneous sequence generation under background noise.** **(a)** Perlin landscape showing preferred directions of projection for excitatory neurons. **(b)** Spontaneously generated sequences visible in spike raster of 14400 excitatory neurons over 2 seconds of background noise. Each point is a spike, every 100th spike is shown. **(c)** Cumulative explained variance for PCA computed on spike rasters for fraction silenced from $p_{inh} = 0$ to $p_{inh} = 0.9$. Dashed line shows 70% variance explained as reference to estimated dimensionality (> 14D, higher dimensional than evoke sequences, see Fig. 5b in main text). Each line is averaged over 5 subspaces. **(d)** Principal angles between different subspaces for fraction silenced from $p_{inh} = 0$ to $p_{inh} = 0.9$. Notice the range of behavior from highly aligned to nearly orthogonal. Each line shows the principal angles between one pair of subspaces with the same fraction silenced, see color code. With 5 subspaces per $p_{inh}$ this gives $\frac{5 \cdot 4}{2} = 10$ comparisons. **(e)** First principal angle as a function of fraction silenced. Panels (b,c,d) correspond to Fig. 5e,f,g in main text, here for spontaneously generated sequences, in main text for evoked sequences.
